# RdDM mutants reduce histone methylation rather than DNA methylation at the paramutated maize *b1* enhancer

**DOI:** 10.1101/2023.08.29.555344

**Authors:** Iris Hövel, Rechien Bader, Marieke Louwers, Max Haring, Kevin Peek, Jonathan I. Gent, Maike Stam

**Affiliations:** Swammerdam Institute for Life Sciences, Universiteit van Amsterdam, P.O. box 1210, 1090 GE Amsterdam, The Netherlands; argenx BV, Industriepark Zwijnaarde 7, 9052 Zwijnaarde (Ghent), Belgium; Universiteit van Amsterdam, P.O. box 19185, 1000 GD Amsterdam, The Netherlands; Department of Plant Biology, University of Georgia, Athens, Georgia 30602, USA

## Abstract

Paramutation is the transfer of mitotically and meiotically heritable silencing information between two alleles. With paramutation at the maize *booster1* (*b1*) locus, the low expressed *B’* epiallele heritably changes the high expressed *B-I* epiallele into *B’* with 100% frequency. This requires specific tandem repeats and multiple components of the RNA-directed DNA methylation (RdDM) pathway, including the RNA-dependent RNA polymerase (RdRP, *mediator of paramutation1*, MOP1), the second-largest subunit of RNA polymerase IV and V (NRP(D/E)2a, *mediator of paramutation2,* MOP2), and the largest subunit of RNA Polymerase IV (NRPD1, *mediator of paramutation3*, MOP3). Mutations in *mop* genes prevent paramutation and release silencing at the *B’* epiallele. In this study we investigated the effect of mutations in *mop1*, *mop2* and *mop3* on chromatin structure and DNA methylation at the *B’* epiallele, and especially the regulatory hepta-repeat 100 kb upstream of the *b1* gene.

We show that mutations in *mop1* and *mop3* result in decreased repressive histone modifications H3K9me2 and H3K27me2 at the hepta-repeat. Associated with this decrease are partial activation of the hepta-repeat enhancer function, formation of a multi-loop structure, and elevated *b1* expression. In *mop2* mutants, which do not show elevated *b1* expression, H3K9me2 and H3K27me2 and a single-loop structure like in wild type *B’* are retained. Surprisingly, high DNA methylation levels at the *B’* hepta-repeat remains in all three mutants. Our results raise the possibility of MOP factors mediating RNA-directed histone methylation rather than RNA-directed DNA methylation at the *b1* locus.

## Introduction

Paramutation involves meiotically heritable epigenetic changes in gene expression caused by *trans*-communication between homologous alleles (Hövel et al., 2015; Hollick, 2017; Springer and McGinnis, 2015; Ronsseray, 2015). Paramutation has been studied in plants, such as maize and tomato, but also in animals, such as mice, Drosophila and *C. elegans*. In maize, paramutation has been best studied at genes encoding transcriptional regulators of the flavonoid pigmentation pathway, *r1 (red1)*, *b1* (*booster1)*, *pl1 (plant color1)*, and *p1* (*pericarp color1*). Most alleles of these genes do not participate in paramutation; they are neutral to paramutation; only specific alleles do. The ability to participate in paramutation is often dependent on repeated sequences (for reviews see Hövel et al., 2015; Hollick, 2017). The alleles participating in paramutation at the *b1* locus are *B’* and *B-I*. *B’* and *B-I* have the same DNA sequence; they are epialleles (Stam et al., 2002a). *B’* is the paramutated, inactive state of *b1* (light pigmented plant), and *B-I* the paramutable, active state of *b1* (dark pigmented plant). In a *B’*/*B-I* heterozygote, the low expressed *B’* epiallele is paramutagenic; it changes the high expressed *B-I* epiallele into *B’* with 100% efficiency, reducing the *B-I* transcription rate 10-20 fold (Patterson et al., 1993). The change of *B-I* into *B’* is mitotically and meiotically heritable and requires multiple tandem copies of part of an 853-nt sequence located ∼100 kb upstream of the *b1* transcription start site (TSS) (Stam et al., 2002a; Stam et al., 2002b; Belele et al., 2013). *B-I* and *B’* carry seven copies of the 853-nt sequence (hepta-repeat), while *b1* alleles insensitive (neutral) to paramutation carry one copy of the repeat unit. Our previous study implicated DNA hypermethylation at the hepta-repeat in both the maintenance and establishment of silencing at the *b1* hepta-repeat, while the presence of Histone H3 lysine K27 dimethylation (H3K27me2) at the hepta-repeat and *b1* coding region was associated with the maintenance of the silenced *B’* state (Haring et al., 2010). Besides their role in paramutation, the 853-nt tandem repeats carry transcriptional enhancer sequences that are required for tissue-specific enhancement of *b1* expression (Belele et al., 2013; Louwers et al., 2009a). In *B-I*, tissue-specific activation of *b1* expression in husk tissue is associated with H3 acetylation (at K9 and K14) and nucleosome depletion at the hepta-repeat (Haring et al., 2010). High *b1* expression levels are in addition associated with chromosomal interactions between the hepta-repeat, TSS and other regulatory regions at the *b1* locus, ∼15, ∼47 and ∼107 kb upstream of the TSS (Louwers et al., 2009a). In *B’*, chromosomal interactions with the TSS, but not with the ∼15, ∼47 and ∼107 kb regions were observed.

To this date, several genes, called *mediator of paramutation* (*mop*) and *required to maintain repression (rmr)*, have been identified to be required to establish paramutation at multiple maize loci (*b1, pl1, r1, p1*) (Dorweiler et al., 2000; Hollick et al., 2005; Sidorenko et al., 2009; Sidorenko and Chandler, 2008; Sloan et al., 2014; Stonaker et al., 2009). In mutants of these genes, the paramutagenic alleles can no longer silence the expression of paramutable alleles. The *mop1* and *rmr* genes identified to play such role in maize share homology with genes involved in the RNA-directed DNA methylation (RdDM) pathway in *Arabidopsis thaliana*. This pathway induces *de novo* DNA methylation (5mC) in all sequence contexts (CG, CHG and CHH, where H is an A, T, or C) at transposable elements and selected genes (Matzke and Mosher, 2014; Rymen et al., 2020; Zemach et al., 2013). In the RdDM pathway, plant-specific RNA Polymerase IV (Pol IV) transcripts are converted into double-stranded RNA by RNA-DEPENDENT RNA POLYMERASE 2 (RDR2), and processed into 24-nt small interfering RNAs (siRNAs) by DICER-LIKE 3 (DCL3). One strand of the siRNAs is incorporated into a complex containing ARGONAUTE 4 (AGO4), targeting the resulting complex to complementary transcripts generated by RNA Polymerase V (Pol V). Subsequently, DOMAINS REARRANGED METHYLTRANSFERASE2 (DRM2) is recruited, mediating *de novo* DNA methylation of cytosines in all sequence contexts (CG, CHG and CHH; H = A, T or C). This in turn leads to recruitment of histone modifying enzymes and chromomethyltransferases such as CROMOMETHYLASE 3 (CMT3), which methylates DNA in the CHG context. It also leads to recruitment of the replication-coupled DNA methyltransferase MET1, which methylates in the CG context.

The first gene identified to be required for the establishment of paramutation was *mop1*, encoding a putative ortholog of RDR2 in Arabidopsis (Alleman et al., 2006; Dorweiler et al., 2000; Haag et al., 2014; Woodhouse et al., 2006a). Subsequently *mop2/rmr7* and *mop3/rmr6* were identified (Erhard et al., 2009; Haag et al., 2014; Hollick et al., 2005; Sidorenko et al., 2009; Sloan et al., 2014; Stonaker et al., 2009); *mop2/rmr7* encodes the maize ortholog of NRP(D/E)2a, the second largest subunit of Pol IV (NRPD1) and Pol V (NRPE1), while *mop3/rmr6*, encodes NRPD1, the largest subunit of Pol IV. Mutations in *mop1*, *mop2/rmr7* and *rmr6* have been shown to cause a severe decrease in the global 24-nt siRNA levels, which is in line with their expected function in the RdDM pathway (Erhard et al., 2009; Jia et al., 2009; Nobuta et al., 2008; Sidorenko et al., 2009; Stonaker et al., 2009). In addition, for *mop1-1* and *mop2-1* mutants, diminished siRNAs levels at the *b1* hepta-repeat were shown (Arteaga-Vazquez et al., 2010; Sidorenko et al., 2009). The nature of the contribution of RdDM is, however, still unclear. The *b1* repeats are transcribed and yield siRNAs not only from *B’*, but also *B-I* and a neutral *b1* allele, suggesting that presence of siRNAs is not sufficient for paramutation (Alleman et al., 2006; Arteaga-Vazquez et al., 2010; Sidorenko et al., 2009).

In line with a role for RdDM, *mop1*, *mop2* and *mop3* mutations are shown to induce a reduction in DNA methylation at specific transposons, especially at DNA sequence regions with high levels of DNA methylation in a CHH context, also called mCHH islands (Gent et al., 2014, 2013; Li et al., 2015, 2014). This reduction in DNA methylation at mCHH islands was indicated to occur in all cytosine contexts and was strongest in *mop3* and weakest in *mop2* mutants. Decreased DNA methylation levels were associated with increased expression of several transposons (Jia et al., 2009; Li et al., 2015; Woodhouse et al., 2006b).

The *mop1*, *mop2/rmr7* and *mop3/rmr6* gene products are not only necessary for paramutation, they are also reported to be required to maintain the silent state of *B’* and *Pl’*, the paramutagenic *b1* and *pl1* loci (Dorweiler et al., 2000; Hollick et al., 2005; Sidorenko et al., 2009; Sloan et al., 2014; Stonaker et al., 2009). In these mutants, increased expression and/or plant pigmentation was observed. Upon crossing out the mutations, depending on the gene and the mutation studied, the epialleles either lose expression and become paramutagenic again or retain expression and are indistinguishable from their non-paramutated states. The first is true for *B’; B’* does not revert to *B-I* in any of the three mutants (Dorweiler et al., 2000) Similarly, *Pl’* does not revert to the active *Pl-Rh* state in *rmr7* mutants (Stonaker et al., 2009). Thus, the epigenetic memory of *B’* and *Pl’* persists in mutants in spite of increased epiallele expression. In *rmr6* mutants, however, the epigenetic memory of *Pl’* can be erased: *Pl’* can stably revert to *Pl-Rh* in *rmr6* mutants (Hollick et al., 2005).

To get a better understanding of the role of the *mop* genes in the maintenance of *B’* silencing, more specifically their role in the maintenance of DNA methylation and repressive histone modifications, we examined the effect of *mop1*, *mop2* and *mop3* mutants on chromatin structure and DNA methylation at the *B’* epiallele, and especially the regulatory hepta-repeat 100 kb upstream of the *b1* coding sequence. Surprisingly, our results show that none of the three *mop* genes is required to maintain high DNA methylation levels at the *B’* hepta-repeat. High DNA methylation levels are maintained in all three mutants. In *mop2* mutants, the *B’* epiallele in addition remains low expressed and retains high levels of the repressive histone modifications H3K9me2 and H3K27me2. The elevated *b1* expression levels in *mop1* and *mop3* mutants are, however, associated with significant reductions in H3K9me2 and H3K27me2 at the hepta-repeat. We propose that this decrease in repressive histone modifications allows the activation of the hepta-repeat enhancer function despite the high DNA methylation levels. Our results indicate that MOP1 and MOP3 play a role in maintaining H3K9me2 and H3K27me2 independent of a role in establishing DNA methylation.

## Results

### Approach

To disentangle the effect of the *mop1-1, mop2-1, mop2-2* and *mop3-1* mutations (Alleman et al., 2006; Dorweiler et al., 2000; Li et al., 2015; Sidorenko et al., 2009; Sloan et al., 2014) on the epigenetic marks and chromatin features that define the mitotically and meiotically heritable *B’* state from the effect on tissue-specific activation of *B’* expression, where relevant, experiments were performed on two types of tissues: seedling and husk tissue. In seedling tissue, comprising 1-month old seedlings with the exposed leaf blades and roots removed, *B’* as well as *B-I* are very low expressed (Haring et al., 2010). In husk tissue (the leaves surrounding the ear) expression of *b1* is transcriptionally activated, resulting in high *B-I* and low *B’* expression*. B’* in a wild-type background was compared to *B’* in *mop* mutant backgrounds. Plants carrying the *B-I* allele served as a positive control for an active *b1* epiallele. The *mop1-1, mop2-2* and *mop3-1* mutations act recessive in preventing paramutation between *B’* and *B-I*, and in their effect on pigmentation (Dorweiler et al., 2000; Sidorenko et al., 2009; Sloan et al., 2014), therefore for these mutants only homozygous mutant tissues were used in our analyses. The *mop2-1* mutation acts dominant in preventing paramutation, and was reported to act recessive for enhancing pigmentation (Sidorenko et al., 2009). Tissues of both heterozygous *Mop2/mop2-1,* and homozygous *mop2-1* mutants were used in our analyses. In the text individuals homozygous for *B’* and a *mop* mutation are referred to as *B’ mop1-1, B’ mop2-1, B’ mop2-2,* and *B’ mop3-1. Mop2 B’*/*mop2-1 B’* individuals are referred to as *B’ Mop2/mop2-1.* Wild type *B’* and *B-I* are referred to as *B’* and *B-I*.

### *B’* expression levels are increased in *mop1* and *mop3*, but not *mop*2 mutants

To monitor the *b1* gene expression levels in the *mop* mutants in our greenhouse conditions, we performed RNA blot analysis using seedling and husk tissue from the mutants as well as *B’* and *B-I* wild type individuals as comparison.

In seedling tissue of *B’ mop1-1, B’ Mop2/mop2-1, B’ mop2-1*, *B’ mop3-1* and *B-I* no expression of *b1* could be detected (triplicate samples; data not shown). As reported previously, in wild-type husk tissue, in which the *b1* gene is transcriptionally activated, the *B’* allele was lowly expressed, while *B-I* transcript levels were increased ∼20-fold compared to the *B’* levels (Figure 1) (Louwers et al., 2009a). In husk tissue from *B’ mop1-1* and *B’ mop3-1*, the *b1* expression levels were ∼3 fold and ∼5-fold increased relative to that in *B’* plants, respectively (Figure 1). The latter upregulation is in line with an 8-fold increase of *B’* transcription in husk tissue observed in an *rmr6-1* mutant (Erhard et al., 2009; Hollick et al., 2005). Both *B’ mop1-1* and *B’ mop3-1* adult plants display as intensely dark purple tissues as *B-I* wildtype plants, indicating that the *b1* RNA expression levels in all three genotypes are above a threshold needed to activate the maize pigmentation pathway. In *B’ Mop2/mop2-1* husk tissue, expression of *B’* remained low, while the expression in *B’ mop2-1* and *B’ mop2-2* was slightly higher, in line with (slightly) higher pigment levels (Figure 1 and Supplemental Figure S1). In conclusion, although all three mutations prevent paramutation between *B’* and *B-I*, the RNA blot data shows they have different effects on the *B’* expression level.

**Figure 1.**
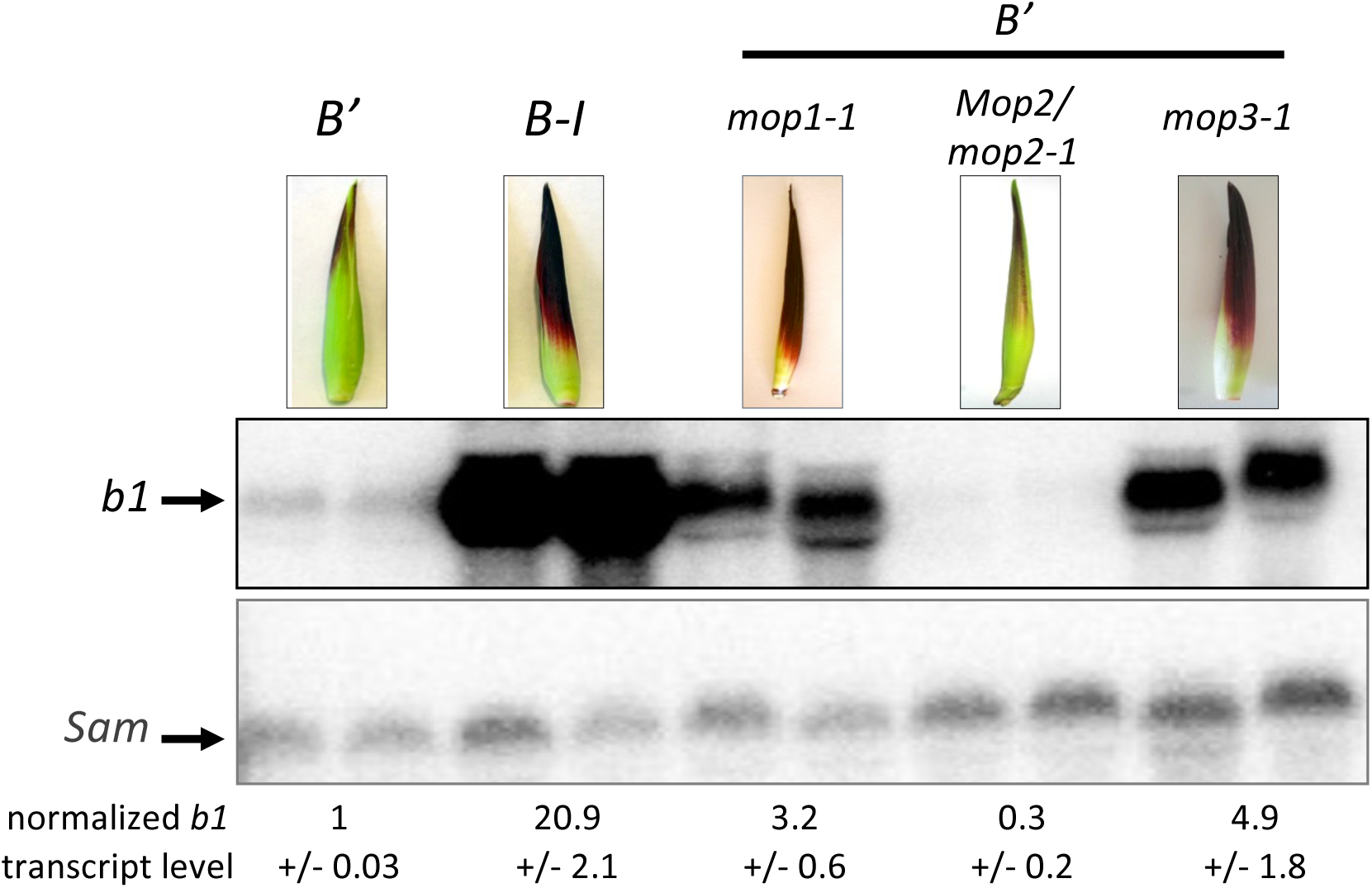
*b1* expression levels in maize husk tissues. RNA blot analysis of RNA from *B’*, *B-I* and *B’ mop1-1, B’ Mop2/mop2-1, B’ mop3-1* husk tissue, using probes recognizing the coding region of *b1* (exons 7-9) and *Sam* (see Supplemental Figure S2B). The intensities of the bands representing full-length transcripts were quantified. The indicated values represent the average plus standard deviation of *b1* transcript levels normalized to *Sam* transcript levels.

### Regulatory sequences at the *b1* locus are activated in *B’ mop1-1* and *B’ mop3-1*

To test whether changes in *b1* transcript level in *B’ mop1-1*, *B’ mop2* and *B’ mop3-1* plants are mediated by the activation of the previously identified distant regulatory sequences at the *b1* locus (Belele et al., 2013; Louwers et al., 2009a; Stam et al., 2002a), we performed chromatin immunoprecipitation (ChIP) with an anti-histone H3K9K14 acetylation (H3ac) antibody on chromatin isolated from husk, tissue in which the *b1* gene is transcriptionally activated. The precipitate was analyzed by quantitative PCR (qPCR) and normalized to *actin1* values (Haring et al., 2007). The targeted genomic regions are shown in Figure 2A, and Supplemental Figure S2; the primers used are listed in Supplemental Table S1. Compared to the levels in *B’* wild-type, elevated levels of H3ac were clearly observed in husk tissue of *B’ mop1-1* and *B’ mop3-1* at the *b1* locus (Figure 2B; Supplemental Figure S3 and Supplemental Table S2), whereby the H3ac levels at the hepta-repeat where generally higher in *B’ mop3-1* than *B’ mop1-1* tissue. This is in line with the higher *B’* expression levels in *mop3-1* compared to *mop1-1*. The H3ac levels in both genotypes were, however, generally still significantly lower than in the highly transcribed *B-I* genotype, indicating that the enhancer sequences at the *b1* hepta-repeat are intermediately activated in *mop1-1* and *mop3-1* plants. In line with low *B’* transcript levels, in heterozygous and homozygous *mop2-1* mutants, *B’* showed only very low H3ac levels at the hepta-repeat and other regions at the *b1* locus, similar as seen for *B’* in a wild-type background (Figure 2B; Supplemental Figure S3 and Supplemental Table S2). In conclusion, the higher *b1* transcript levels in *B’ mop1-1* and *B’ mop3-1* husk tissue are associated with higher histone acetylation levels at the hepta-repeat, indicating activation of the distant regulatory sequences at the *b1* locus.

**Figure 2.**
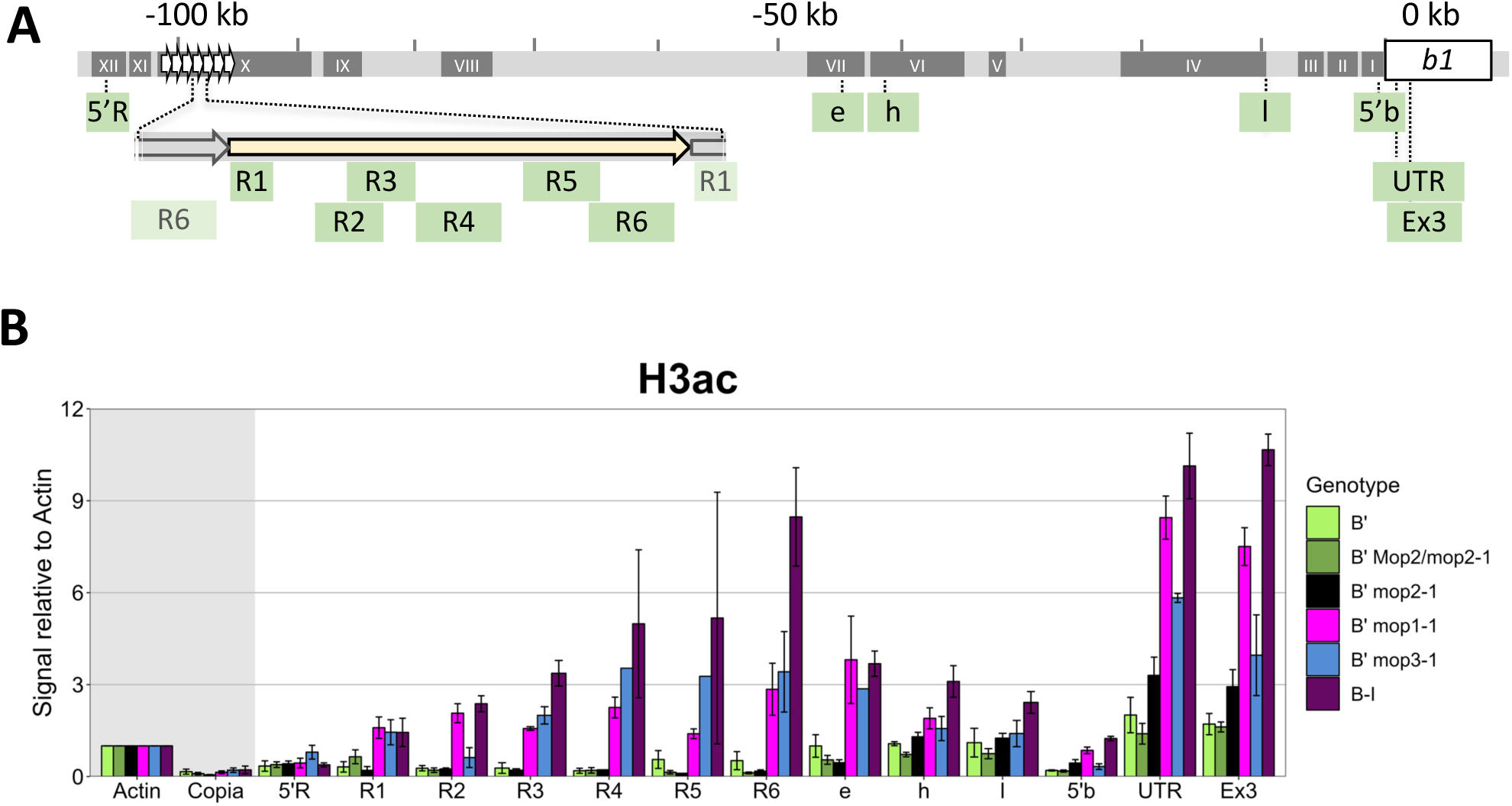
The regulatory hepta-repeat is activated in *mop1-1* and *mop3-1*, but not in *mop2* mutants. A) Schematic representation of the *b1* locus including coding region (white box) and hepta-repeat enhancer (arrowheads). Regions monitored in ChIP are indicated below in green. The fragments (I-XII) examined by 3C (see Figure 3) are indicated as dark grey boxes. B) ChIP-qPCR experiments were performed on husk tissue from *B’*, *B’ mop1-1*, *B’ Mop*/*mop2-1*, *B’ mop2-1*, *B’ mop3-1* and *B-I* plants with an antibody recognizing H3ac. ChIP signals were normalized to *actin* ChIP signals. Error bars indicate standard error of the mean (SEM) of three *(B’*, *B’ mop1-1*, *B’ Mop*/*mop2-1)*, or four (*B’ mop2-1*, *B’ mop3-I, B-I)* biological replicates. See Supplemental Table S2 for summary of ChIP statistics.

### Activated *b1* regulatory sequences are associated with a multi-loop structure

The low expression level of the *B’* epiallele in husk tissue has been associated with the formation of a single loop between the *b1* TSS and hepta-repeat 100 kb upstream, whereas transcriptional activation of the high expressed *B-I* epiallele has been associated with the formation of a multi-loop structure between the TSS and regulatory regions ∼15, ∼47, ∼100 and ∼107 kb upstream (Louwers et al., 2009a). To examine if the activation of the regulatory sequences at the *b1* locus in the *mop* mutants is associated with the formation of a multi-loop structure, chromosome conformation capture (3C) was applied, using husk tissue from *B-I*, *B’* (Louwers et al., 2009a) and *B’ mop1-1*, *B’ Mop2/mop2-1* and *B’ mop3-1* plants, and *Bgl*II as restriction enzyme. 3C allows the identification of physical interactions between selected genomic regions. The TSS (fragment 1), hepta-repeat (fragment X) and a fragment ∼47 kb upstream (fragment VII) were used as a viewpoint (bait) (Figure 3, Supplemental Figure S2). The *S-adenosyl methionine decarboxylase* (*Sam*) locus was used as an unrelated, internal control for data normalization (Louwers et al., 2009a).

**Figure 3.**
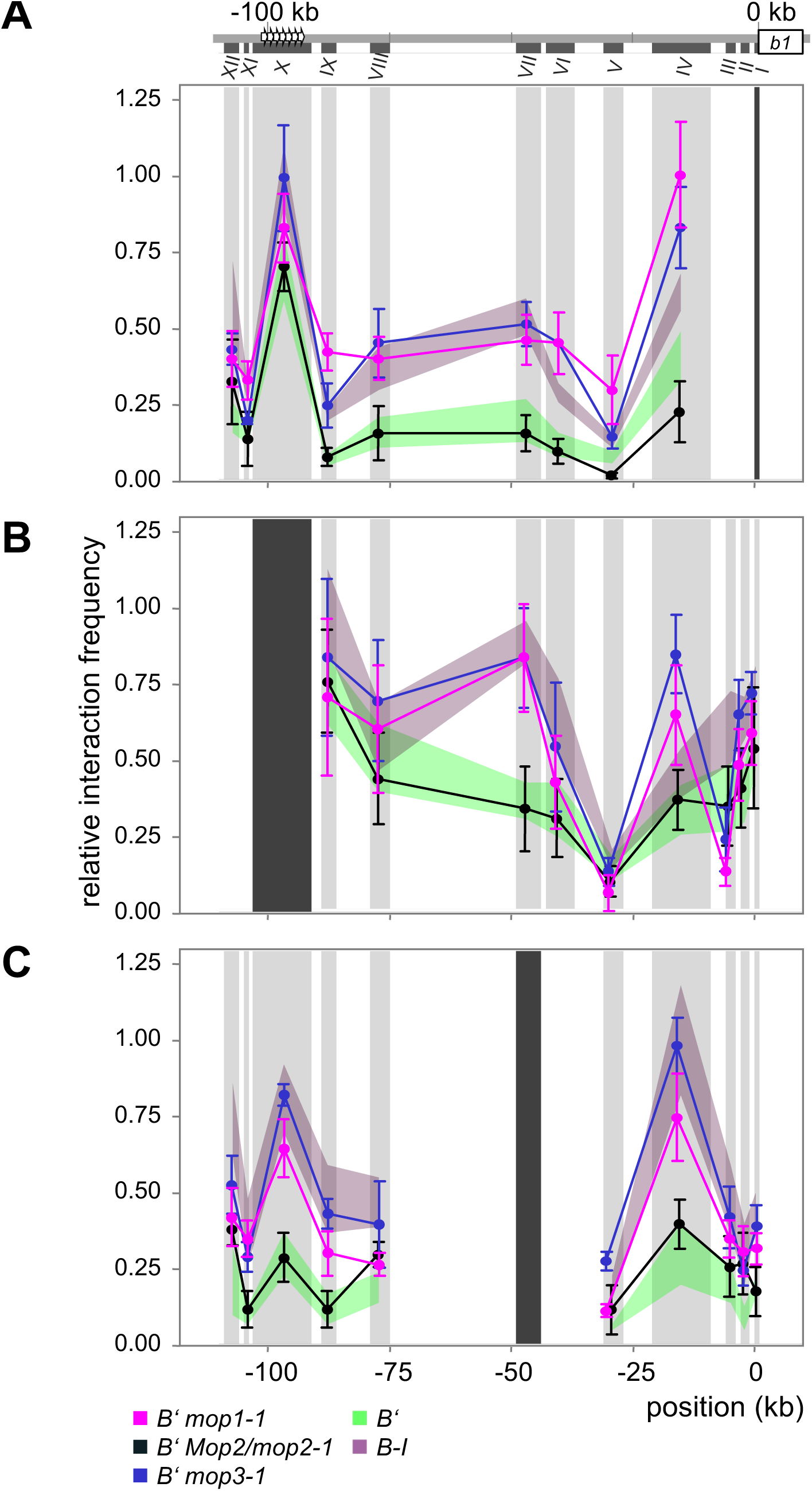
*B’* shows a multi-loop structure in the *mop1-1* and *mop3-1* mutant background, and a single loop structure in *Mop2/mop2-1* mutant background. At the top a schematic representation of the *b1* locus including the coding region (white box) and hepta-repeat enhancer (arrowheads). The *Bgl*II fragments (I-XII) examined by 3C are indicated as dark grey boxes. The viewpoints I (TSS; panel A), X (Hepta-repeat; panel B), and VII (∼47kb upstream; panel C) are indicated by black vertical bars. Data were normalized using crosslinking frequencies measured for the *Sam* locus (Supplementary Figure S2). Error bars indicate the SEM of four biological replicates each for *B’ mop1-1*, *B’ Mop2/mop2-1* and *B’ mop3-1*. Data for *B’* and *B-I* (reproduced from Louwers et al., 2009) are shown in shading for comparison. The experiments for *B’*, *B-I*, and the *mop* mutants were done in parallel.

Our data show that in *mop1-1* and *mop3-1* husk tissue, the *B’* epiallele shows a *B-I*-like multi-loop conformation, whereas in *Mop2/mop2-1* tissue *B’* shows a *B’*-like single-loop conformation (Figure 3A-C). When using fragment I (*b1* TSS) as viewpoint, in all samples the hepta-repeat (fragment X) showed high interaction frequencies (Figure 3A), whereby the *B’* epiallele showed the lowest interaction frequencies in a wild-type and *Mop2/mop2-1* background, and the highest in a *B’ mop3-1* and *B-I* background. In addition, fragment I also interacted with fragments XII (−107 kb), VII (−47 kb), and VI (−40 kb) and IV (−15 kb), whereby high interaction frequencies with fragments VII, VI and IV where only observed in *B’ mop1-1*, *B’ mop3-1* and *B-I* tissue, consistent with a multi-loop structure. When using fragment X and VII as viewpoints (Figure 3, B and C), the high interaction frequencies between the TSS, hepta-repeat and regions ∼47, and ∼15 kb upstream were confirmed for *B’ mop1-1* and *B’ mop3-1*. The interactions involving the region ∼107 kb upstream were confirmed for all genotypes, with *B’* in a wild type and *Mop2/mop2-1* background showing the lowest interaction frequencies.

In summary, our 3C data show that the transcriptional enhancement of *B’* expression in the *mop1-1* and *mop3-1* mutant background is associated with the formation of a *B-I*-like multi-loop structure, whereas the low *B’* expression level in the *Mop2/mop2-1* background is associated with a *B’*-like single-loop structure as observed.

### *B’* hepta-repeat DNA methylation level is retained in all *mop* mutants

At RdDM-target loci, the *mop1, mop2* and *mop3* genes have been indicated to play a major role in maintaining CHH methylation (mCHH), and a minor role in maintaining CG and CHG methylation (mCG, mCHG) (Gent et al., 2014; Li et al., 2015, 2014). In these studies, a decrease in mCG and mCHG seemed most prominent in a *mop3* and least prominent in a *mop2* mutant. Previous experiments showed that the silenced *B’* allele is DNA hypermethylated at the hepta-repeat junction regions compared to *B-I* (Haring et al., 2010). To examine if transcriptional activation of *B’* in *mop* mutants is associated with a decrease in DNA methylation, DNA blotting and bisulfite sequencing experiments were performed.

DNA blot analyses were performed as described in Haring *et al*. (2010), using genomic DNA from leaf and husk tissue, isolated from 4 to 27 individuals per genotype at different stages of development. DNA was isolated and digested with *Eco*RI or *Bam*HI, which released a discrete fragment containing the entire hepta-repeat, and eight different methylation sensitive restriction enzymes (Figure 4), size fractionated, blotted and hybridized with a repeat probe (for representative examples, and the sequence and sites examined see Supplemental Figure S2, S4 and S5). To check for complete digestion, probes recognizing unmethylated sequence regions within the *b1* locus were used. To approximate the DNA methylation levels, relative band intensities of all detected restriction fragments were computationally compared to all theoretical possible combinations of fragment intensities.

**Figure 4.**
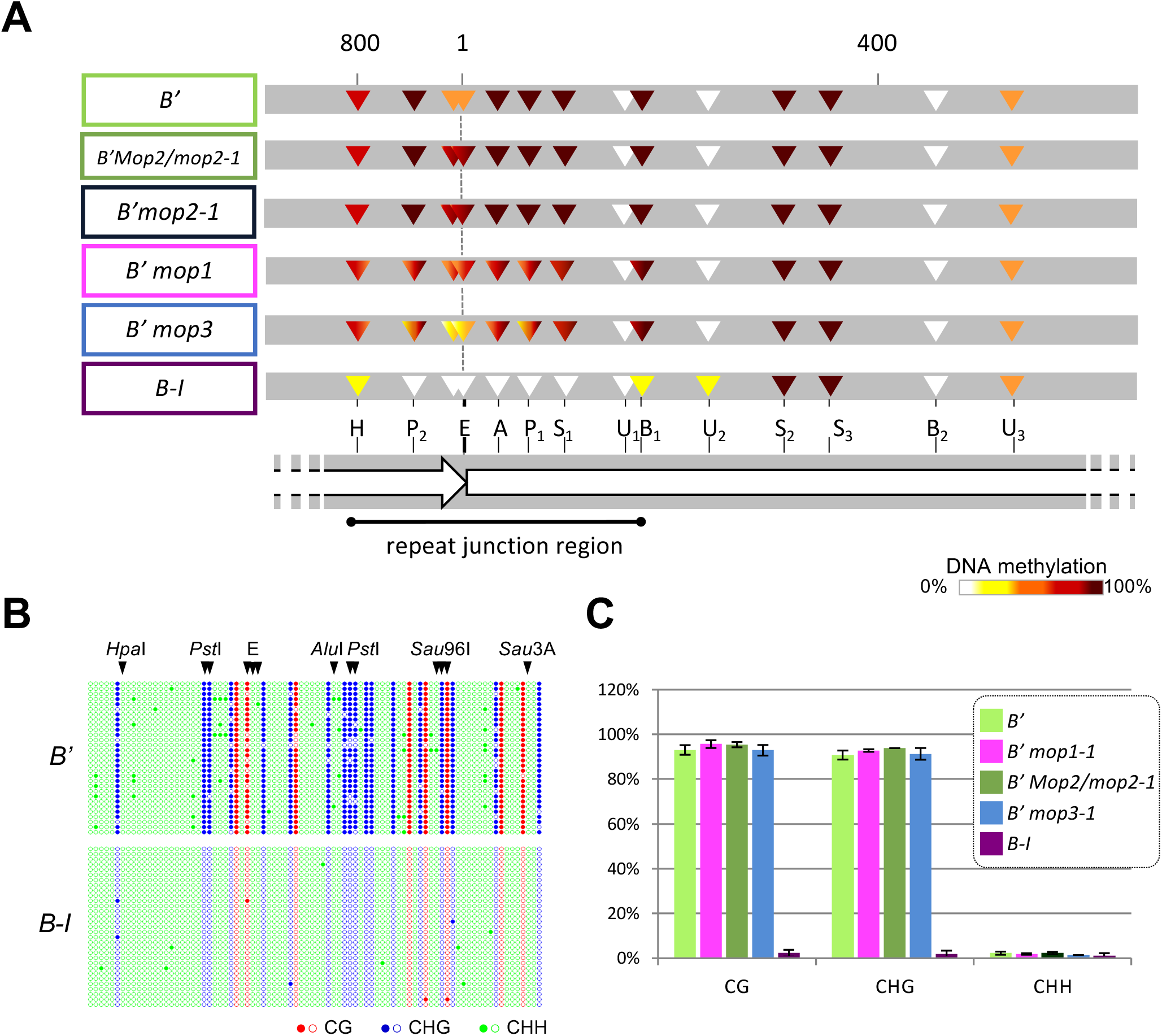
Differentially methylated repeat junction region retains high levels of symmetric DNA methylation in *mop* mutants. A) Summary of the DNA methylation levels data obtained by DNA blot analysis. DNA derived from *B’* wild type, *B-I* and *B’ mop* mutants was digested with the methylation-insensitive *Eco*RI or *Bam*HI and the methylation-sensitive enzymes indicated. A zoom of part of the hepta-repeat, spanning the repeat junction region, is shown. The consensus repeat methylation profiles of *B’* and *B-I* (Haring et al., 2010) are shown. DNA methylation levels are specified by colors: 0-12.5%, white; 12.5-37.5%, yellow; 37.5-62.5%, orange; 62.5-87.5%, red; 87.5-100% methylation, dark red. H, *Hpa*I; P, *Pst*I; E, *Hha*I and *Hae*II; A, *Alu*I; S, *Sau*96I; U, *Sau*3AI; B, *Bsm*AI; Numbers discriminate individual sites present more than once every repeat. For details on the quantification of the DNA methylation levels see Haring *et al*. (2010). B) DNA methylation profiles of the *b1* repeat junction region at base-pair resolution for *B’* and *B-I*, generated by targeted bisulfite sequencing. This region includes the R1 and R6 loci assayed by ChIP-qPCR. See Supplementary Fig. S8 for all data sets. The dot-plots show the DNA methylation for 30 and 31 individual *B’* and *B-I* clones, respectively. Methylated and unmethylated cytosines are represented by filled and empty circles, respectively. Red, CG; Blue, CHG; Green, CHH. The restriction sites examined by DNA blotting are indicated above the panel for comparison. E indicates a site cut by both *Hha*I and *Hae*II. C) Total fraction of methylated cytosines in the repeat junction region in all three sequence contexts as measured by bisulfite sequencing. The mean and standard deviations shown are calculated from three (*B’ mop1-1*) or two (all other genotypes) biological replicates. *M/mop2* indicates *Mop2/mop2-1*.

Compared to *B’* tissue, in *B’ mop1-1* and *B’ mop3-1* tissue we observed only a slight decrease in DNA methylation level at the *B’* repeat junction regions, with the most decrease observed in *B’ mop3-1* tissues (Figure 4, Supplemental Figure S4 and S6). The decrease occurred in several, but not all plants, and was observed at specific restriction sites, in a stochastic manner; a decrease at a particular site was not necessarily associated with a decrease at another site in the same sample. In *B’ Mop2/mop2-1, mop2-1* as well as *mop2-2* samples, no decrease in DNA methylation was observed at the hepta-repeat (Figure 4, Supplemental Figure S4 and S6).

In addition to the slight decrease in DNA methylation, in most *B’ mop1-1* and all *B’ mop2* samples we detected hypermethylation at the repeat junctions (Figure 4 and Supplemental Figure S4 and S6; *Hha*I and *Hae*II). In the *mop3-1* mutant, however, a stochastic decrease in DNA methylation was observed at the site hypermethylated in the *mop1-1* and *mop2* mutants.

We expected that during the development of *B’ mop1-1* plants, as previously hypothesized (Ohtsu et al., 2007), a progressive decrease in DNA methylation levels would be observed due to cell division. To test this hypothesis, DNA blot analysis was performed on DNA from six to eight leaves collected during the development of six homozygous *B’ mop1-1* plants (Supplemental Figure S7). In most plants monitored (5 of 6), we did detect a slight decrease in DNA methylation, but in general, the maximum decrease in DNA methylation was already observed in the second leaf stage, suggesting that this reduction in DNA methylation may already have occurred in very early developmental stages such as embryos, in which *Mop1* appears higher expressed compared to seedlings and leaves (Sekhon et al., 2011; Woodhouse et al., 2006b).

DNA methylation at the *B’* repeat junction regions in *mop1-1*, *Mop2/mop2-1* and *mop3-1*, and *B’* and *B-I* wild type was also monitored by targeted bisulfite sequencing, which provides single base-pair resolution data on cytosine methylation. DNA was obtained from leaf 4 tissue of V4 stage plants, and *Fie2* was used as a control for complete conversion (Gutiérrez-Marcos et al., 2006). As observed with DNA blot analysis, the *B’* repeat junction region was hypermethylated compared to that of *B-I*, both in wild-type and *mop* mutant leaf tissue (Figure 4B, Supplemental Figure S8). Bisulfite sequencing confirmed the lack of significant DNA demethylation in the *mop* mutants. In fact, there was no detectable decrease in DNA methylation. This difference from the DNA blot analysis method could be due to the limited number of samples that can be easily examined by targeted bisulfite sequencing. Also note that bisulfite sequencing represents individual molecules in individual cells while DNA blot analysis represents large population of cells. In addition, there may be variation between the seven repeats, and DNA blot analyses provide the overall picture. Intriguingly, in contrast to the expectations (Li et al., 2014), the vast majority of DNA methylation observed at the *B’* allele was in the CG and CHG context, and the low level in the CHH context (3%) was similar in *B-I* and *B’ mop* samples (Figure 4B and C, Supplemental Figure S8). These findings are in contrast to reported observations that the RdDM pathway, in which the MOP proteins play a role, mediates *de novo* DNA methylation in all sequence contexts, including CHH (Gent et al., 2014; Li et al., 2015, 2014; Matzke and Mosher, 2014).

Together, our data indicate that i) activation of the enhancer function of the hepta-repeat occurs in the presence of high DNA methylation levels, ii) the MOP proteins plays a minor role in maintaining DNA methylation at the *b1* hepta-repeat, and iii) the *B’* repeat junction regions show very low mCHH levels.

### Genome-wide variation in mCHH levels exists at RdDM loci

Our finding that the *B’* repeat junction region, which is likely to be an RdDM locus, has little mCHH, prompted us to explore whether similar loci, with MOP1-dependent 24-nt siRNAs, but little mCHH, are present elsewhere in the genome. To do this, we used published small RNA and DNA methylation datasets from wild-type and *mop1-1* mutant developing ear (Gent et al., 2014). First, we compared the abundance of perfect-matching, uniquely-mapping 24-nt siRNAs across the maize B73 genome to identify loci with abundant siRNAs in wild-type and at least ten-fold reduced coverage in the *mop1-1* mutant. We then categorized all identified loci based on mCHH levels in developing ear, whereby loci with mCHH values of less than 0.02 (mC/total C) were categorized as low mCHH *mop1* loci, and greater than or equal to 0.02 were categorized as high mCHH *mop1* loci. For increased stringency, we used 50-bp non-overlapping intervals across the genome, and only included loci where at least two adjacent loci were both low or both high mCHH. All adjacent 50-bp loci fitting the same category were then merged, producing 69 low mCHH *mop1* loci and 3107 high mCHH *mop1* loci, all of which were at least 100-bp in length (Supplemental Figure S9). Note that these numbers are certainly smaller than the actual numbers of such loci in the genome. For this analysis, our aim was to determine whether the *b1* enhancer is an anomaly, not to rigorously identify and categorize all *mop1* loci in the genome. Excluding multi-mapping siRNAs is the largest limiting factor of this analysis, but was necessary for robust interpretation of the data. Based on these numbers, approximately 2% of *mop1* loci have low mCHH levels, like observed for the *B’* repeat junction region. While both the low and high mCHH *mop1* loci had high abundance of 24-nt siRNAs, the siRNAs at high mCHH loci were on average about two-fold more abundant than at low mCHH loci. Both sets were dominated by 24nt siRNAs, though other lengths were also present and were also similarly more abundant at high mCHH loci (Supplemental Figure S9).

In whole-genome studies like this, selecting for the tails of a distribution (in this case low mCHH) will always yield some number of false positives. To evaluate whether the loci we identified have reproducibly low mCHH, we used data from a different tissue (developing second leaf), produced using a different method (enzymatic conversion rather than bisulfite conversion) (Hufford et al., 2021). In spite of a general reduction in mCHH in leaf compared to ear, the general patterns were highly reproducible, not just for mCHH, but also mCG and mCHG (**Figure 5**). In particular, the mean mCHH in leaf was 0.02 in low mCHH loci, but 0.10 in high mCHH loci, and both mCG and mCHG showed high, heterochromatin-like levels in both sets of loci (Gent et al., 2014). These data also indicate that the majority of low mCHH loci with MOP1-dependent siRNAs in ear also have low mCHH in leaf.

**Figure 5.**
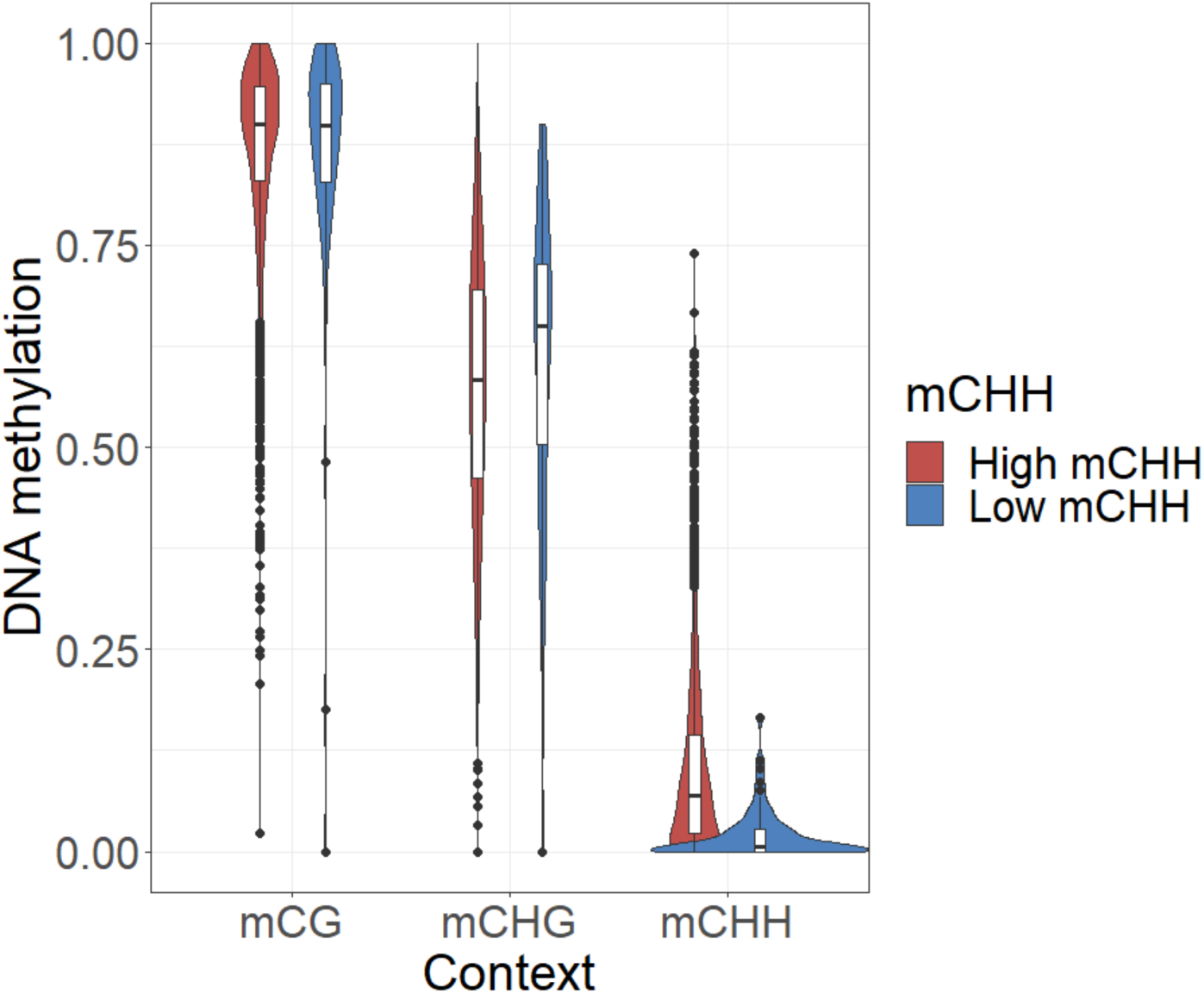
The maize genome contains low-mCHH and high-mCHH RdDM loci. Average DNA methylation levels for high and low mCHH loci in the developing second leaf of B73. Values of 5-methylcytosine are relative to the total number of cytosines (mC/total C) in each sequence context (CG, CHG, or CHH).

### Transcriptional activation is not associated with a decrease in nucleosome occupancy at the *B’* repeats

Nucleosome occupancy plays an important role in gene regulation (Jiang and Pugh, 2009; Struhl and Segal, 2013), with nucleosome depletion at regulatory regions being associated with gene expression. To test whether the increased *B’* expression level in *mop1-1* and *mop3-1* plants (Figure 1) is associated with decreased nucleosome occupancy, ChIP-qPCR experiments were performed using an anti-histone H3 antibody (H3core) that recognizes canonical H3 and H3 variants. As observed before, the hepta-repeats display a low and high nucleosome occupancy in *B-I* and *B’* husk, respectively (Figure 6, Supplemental Figure S10) (Haring et al., 2010). This is consistent with the hepta-repeat acting as a transcriptional enhancer in *B-I*, but not *B’* plants. In husk tissue of *B’ mop1-1* plants, however, compared to *B*’ wildtype plants no significant differences in nucleosome occupancy level were observed at the hepta-repeat; they were significantly different from those in *B-I* plants, but not *B’* plants (Figure 6, Supplemental Figure S10, Supplemental Table S2). This indicates that in *B’ mop1-1*, the increased *B’* RNA level is not facilitated by nucleosome displacement. The histone H3 levels at the hepta-repeat in *B’ mop3-1* plants were not significant different from both *B*’ and *B-I* plants, indicating they were at an intermediate level and the elevated *B’* RNA levels in *B’ mop3-1* plants could be slightly facilitated by nucleosome displacement. *Mop2/mop2-1* plants showed similar results at the *B’* hepta-repeat as *mop1-1* plants, while *mop2-1* homozygous plants showed H3core levels that were significantly different from both *B’* and *B-I* wildtype. They were higher than in *B’*.

**Figure 6.**
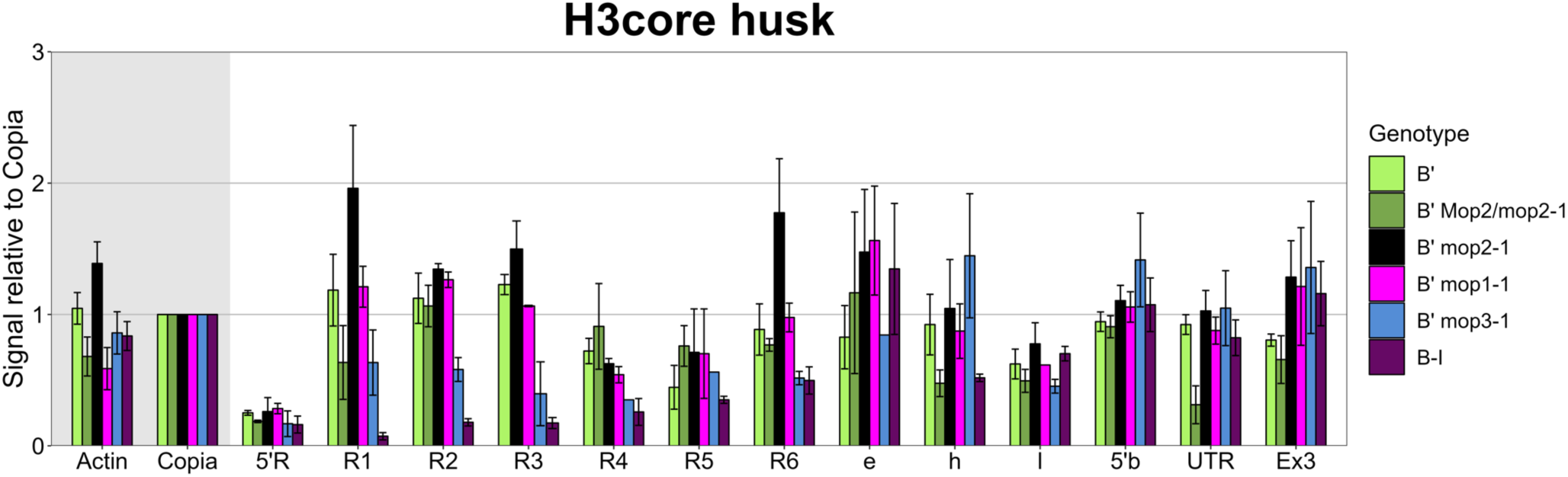
No significant decrease in nucleosome occupancy was observed in the *mop* mutants. ChIP experiments were performed with an antibody recognizing histone H3, using husk tissue from *B’* (light green), *B’ Mop*/*mop2-1* (green), *B’ mop2-1* (black), *B’ mop1-1* (magenta), *B’ mop3-1* (blue), and *B-I* (purple) plants. Data was normalized to *actin* values. The error bars indicate the standard error of the mean (SEM) of two (*B’ mop1-1*), three (*B’*, *B’ Mop*/*mop2-1*, *B’ mop3-1)*, or four (*B’ mop2-1*, *B-I*) replicate experiments. For the no-antibody control signals see Supplemental Figure S10.

### Transcriptional activation is associated with reduced H3K9me2 and H3K27me2 levels

The silencing of transposons and other repeats is associated with DNA methylation, H3K9me2 and H3K27me2 (Bernatavichute et al., 2008; Gent et al., 2014; Liu et al., 2021; Roudier et al., 2011; West et al., 2014). In maize, H3K9me2 and H3K27me2 are both enriched in heterochromatin, and they are enriched at low and moderately expressed genes, respectively (relative to highly expressed genes) (Gent et al., 2014). Our previous study indicated that the *B’* gene body is marked by H3K27me2 in both seedling and husk tissue, while the *B’* hepta-repeat carries H3K9me2 and H3K27me2 only in husk (Haring et al., 2010). To examine if activation of the *B’* epiallele in *mop* mutant plants is associated with reduced H3K9me2 and H3K27me2 levels, ChIP-qPCR experiments were performed on seedling and husk tissue, using monoclonal antibodies with a much better signal-to-noise ratio than the antibodies used in our previous study (Figure 7, see Methods) (Haring et al., 2010). The *B’* and *B-I* epialleles are transcriptionally activated in husk, but not in seedling tissue. A high-copy *Copia* sequence was used to normalize the data (Haring et al., 2010, 2007).

**Figure 7.**
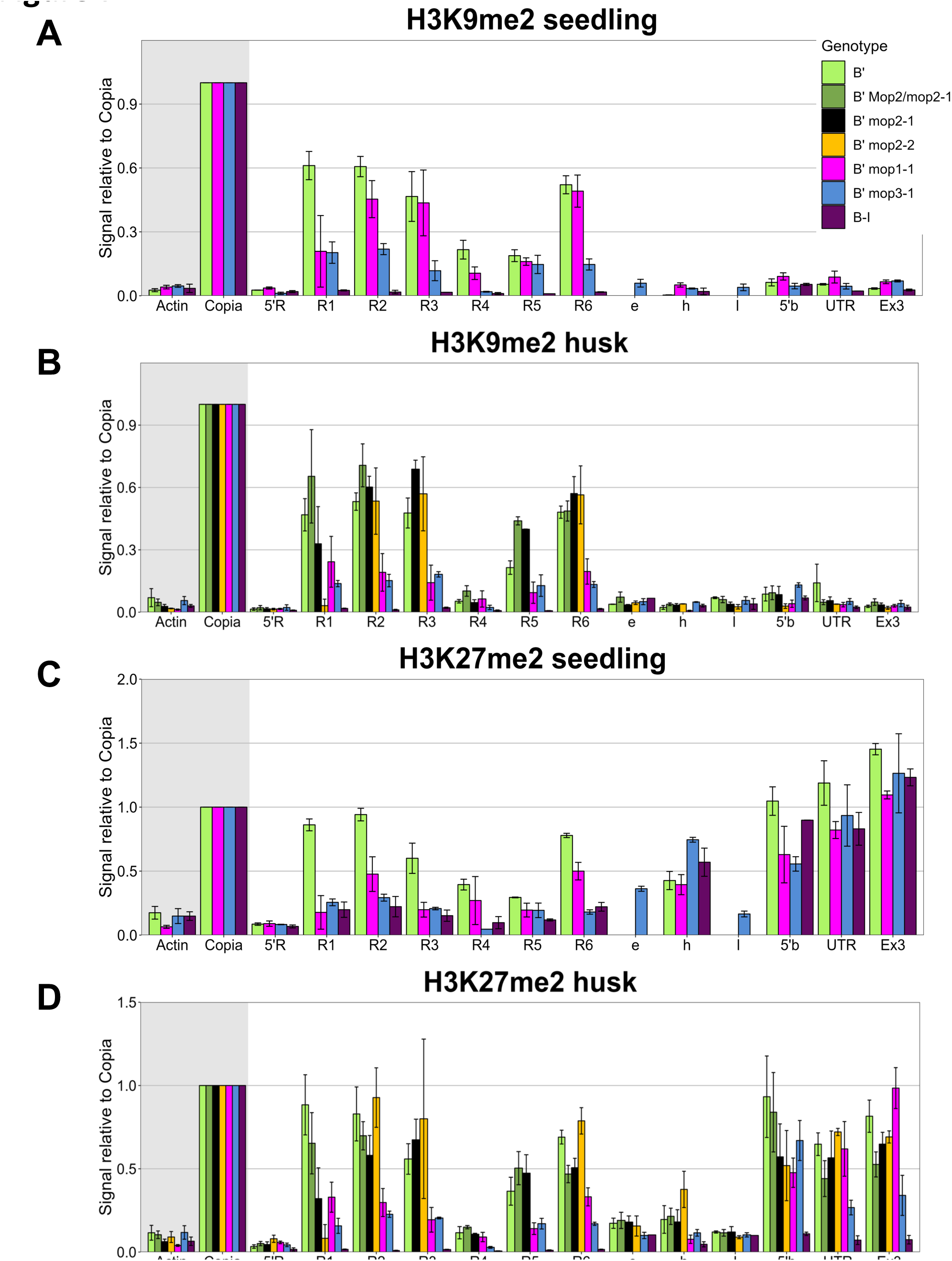
Transcriptional activation in *mop1-1* and *mop3-1* is associated with reduced levels of repressive histone marks. ChIP-qPCR experiments were performed on seedling (A, C) and husk tissue (B, D) from *B’*, *B’ Mop*/*mop2-1*, *B’ mop2-1*, *B’ mop1-1*, *B’ mop3-1* and *B-I* plants with antibodies recognizing H3K9me2 (A, B), and H3K27me2 (C, D). ChIP signals were normalized to *Copia* signals. Error bars indicate the SEM of multiple biological replicates. For seedlings, three (*B’*, *B’ mop1-1*) or two (*B’ mop3-1, B-I)* replicates were performed. For husks (B, D), two (*B’ mop2-2*), three (*B’, B’ mop3-1*, *B-I*), four (*B’ mop1-1, B’ Mop*/*mop2-1*, *B’ mop2-1)* replicates were performed. For the no-antibody control signals see Supplemental Figure S11.

Unlike our previous results, H3K9me2 was significantly enriched at the hepta-repeat in *B’* seedling as well as husk tissue, suggesting that the *B’* hepta-repeat is marked by H3K9me2 in a tissue-independent manner (Figure 7A, B, Supplemental Figure S11A, B). The *b1* gene itself in the *B’* epiallelic state did not show significant H3K9me2 levels, nor in the *B-I* epiallele. In line with H2K27me2 being associated with repressed sequences, H3K27me2 levels were significant enriched at the hepta-repeat, gene body and other sequences in *B’* and *B-I* seedlings, and *B’* husk tissue, tissue in which the *b1* gene is not activated (Figure 7C, D, Supplemental Figure S11C, D, Supplementary Table S2). In *B-I* husk tissue, where *b1* transcript levels are high (Figure 1), only background levels of H3K27me2 were detected across the *b1* locus.

In seedling tissue, in which the *b1* gene is transcriptionally inactive, the H3K9me2 levels were significantly lower at the *B’* hepta-repeat in *mop3-1* plants than in wild type *B’* plants (Figure 7A, C, Supplemental Table S2). The same is true for the H3K27me2 levels for both *mop1-1* and *mop3-1* seedlings. In husk tissue, both the H3K9me2 and H3K27me2 levels were significantly lower in *mop1-1* and *mop3-1* plants compared to wild type *B’* plants (Figure 7B, D and Supplemental Figure S11B, D, Supplementary Table S2). The significant decrease in H3K9me2 (*mop3-1*) and H3K27me2 (*mop1-1* and *mop3-1*) levels at the hepta-repeat is already observed in seedling tissue and therefore independent of transcriptional activation of the *b1* gene, suggesting this decrease is a true effect of these *mop* mutations. At the gene body (UTR, Ex3), only the H3K27me2 levels in *B’ mop3-1* husk tissue were significantly different from those in wild-type *B’* plants, in line with the relatively high expression level in *B’ mop3-1* plants. The increased *b1* expression levels in *B’ mop1-1* husk tissue occurs without a significant decrease of H3K27me2 levels at the gene body.

In *Mop2/mop2-1*, *mop2-1* husk tissue, similar levels of H3K9me2 and H3K27me2 were observed at the *B’* hepta-repeat as in wild type *B’* tissue (Figure 7B, D and Supplemental Figure S11B, D, Supplementary Table S2). H3K27me2 levels at the *b1* coding region were also not significantly different. These observations were confirmed using the independent *mop2-2* mutant, showing that the results are not specific for a dominant mutation like *mop2-1;* repressive chromatin marks at the *B’* epiallele are also retained in the *mop2-2* mutant.

In conclusion, in *mop1-1* and *mop3-1* mutants the repressive H3K9me2 and/or H3K27me2 marks are reduced at the *B’* hepta-repeat. For H3K9me2, this reduction is tissue-independent for *mop3-1*, while for H3K27me2 this is true for both mutants. We propose that these lower levels allow transcriptional activation of the *B’* epiallele in husk tissue. The retention of these marks in *mop2* mutants correlates with the lack of a significant increase in *B’* transcript levels in these mutants.

## Discussion

In this study we show that the RNA-dependent RNA polymerase (RdRP, MOP1) and the largest subunit of RNA polymerase IV (NRPD1, MOP3), proteins in the RNA-directed DNA methylation (RdDM) pathway, are involved in maintaining high levels of the repressive histone marks H3K9me2 and H3K27me2 at the *b1* regulatory hepta-repeat, but not in maintaining high DNA methylation levels. In the *mop1* and *mop3* mutants strong and reproducible reductions in the levels of the repressive H3K9me2 and H3K27me2 histone marks are detected at the regulatory hepta-repeat enhancer, but only weak and stochastic reductions in DNA methylation. We also show that *mop1* and *mop3* mutants, which prevent paramutation at multiple maize loci including *b1*, allow partial activation of the regulatory hepta-repeat enhancer at the *B’* epiallele in the presence of high DNA methylation and nucleosome occupancy levels.

### Role of MOP proteins in maintenance of DNA methylation

Mutations in *Mop1* (RdRP), *Mop2* (NRP(D/E)2a), and *Mop3/Rmr6* (NRPD1) prevent paramutation and significantly reduce overall 24-nt siRNAs levels (Arteaga-Vazquez et al., 2010; Erhard et al., 2009; Nobuta et al., 2008; Sidorenko et al., 2009; Stonaker et al., 2009). The *mop1-1* and *mop2-1* mutants have in addition been shown to significantly reduce the levels of hepta-repeat-derived 24-nt siRNAs (Arteaga-Vazquez et al., 2010; Sidorenko et al., 2009). We, however, did not observe significant reductions in mCG and mCHG at the repeat junction region in any of the mutants examined. Earlier studies in maize did report reductions in DNA methylation at so-called mCHH islands, whereby decreases were more severe in *mop3-1* than *mop1-1* plants and more in mCHG than in mCG (Gent et al., 2014; Li et al., 2014). In addition, studies in Arabidopsis, showed decreased mCHG and mCG levels in *rdr2*, and *nrpd1* mutants (Tang et al., 2016; Zemach et al., 2013). We only observed stochastic decreases in DNA methylation with our DNA blot analyses in *mop1-1* and *mop3-1* mutants, and to a larger extent in *mop3-1* than *mop1-1* plants. Not all plants, however, showed such decrease at the restriction sites monitored, therefore, the activation of the hepta-repeat enhancer in these mutants is only weakly associated with non-reproducible decreases in DNA methylation.

### Role for RdDM components in maintenance of repressive histone modifications

The activation of the enhancer at the *B’* hepta-repeat in the *mop1-1* and *mop3-1* mutants is associated with significant reductions in the repressive H3K9me2 and/or H3K27me2 marks. Given the well-established roles for siRNAs in directing DNA methylation, the reductions in these histone modifications would normally be explained as a downstream consequence of loss of DNA methylation. In principle, the MOP siRNA machinery could also directly target H3K9me2 or H3K27me2 through unidentified protein-protein interactions between argonaute proteins and histone methyltransferases. Such a scenario is supported by the existence of RNA-directed histone modification without RNA-directed DNA methylation in diverse fungi and animals, including *Drosophila melanogaster*, *Caenorhabditis elegans* and *Schizosaccharomyces pombe* (Castel and Martienssen, 2013; Gu et al., 2012; Pal-Bhadra et al., 2004; Verdel et al., 2004). In addition, a few studies have implicated siRNAs in directing H3K9me2 in Arabidopsis (Jackel et al., 2016; Parent et al., 2021), although effects on H3K9me2 and DNA methylation could not be unequivocally disentangled.

Interestingly, it has been shown that tandem repeats are efficiently and stably silenced by H3K9 methylation through RNA-mediated mechanisms (Grewal, 2023). Small RNAs derived from only one or a limited number of repeats can target all the repeats at a given locus and mediate a high density of H3K9 methylation, which in turn recruits histone deacetylases (HDACs) that stabilize heterochromatic regions (Grewal, 2023; Zofall et al., 2022). Similarly, small RNAs may be required to target high H3K9me2 levels at the hepta-repeat, which in turn recruits HDACs stabilizing the silencing.

Resolving the hierarchy of histone methylation and DNA methylation is complicated by indirect effects of developmental gene regulation. In husk, where the hepta-repeat serves as a weak enhancer for the *b1* gene in the mutants, we expect that some of the reduction in histone methylation is due to the presence of transcription factors and associated chromatin modifiers. In seedling tissue, however, in which *b1* is transcriptionally repressed, the reduction should be independent of transcriptional activation of *b1*. Thus, the stronger reduction of H3K27me2 over H3K9me2 in *B’ mop1-1* seedling could suggest a more direct effect of the MOP siRNA pathway on H3K27me2.

A clear link between H3K27me2 and RdDM in plants has not been shown before. H3K27me2 is, as H3K27me3, intimately associated with transcriptional silencing through Polycomb group (PcG) proteins (Conway et al., 2015). While H3K27me3 is mainly associated with repressed genes, H3K27me2 is localized more ubiquitously throughout the genome. Data from animals indicate that, unlike H3K27me3, H3K27me2 is not associated with stable binding of Polycomb Repressive Complex 2 (PRC2); it is indicated to act as a protective layer against unspecific transcriptional activation (Ferrari et al., 2014; Lee et al., 2015). This includes preventing the activation of enhancer sequences in inappropriate cell types. In agreement with these findings, in Arabidopsis and maize, H3K27me2 is enriched at silenced promoters, genes and repeats (Gent et al., 2014; Locatelli et al., 2009; Mathieu et al., 2005; Park et al., 2012; Roudier et al., 2011), and in wheat at euchromatic transposons at distal chromosome ends (Liu et al., 2021). In Arabidopsis, H3K27me2 enrichment seems independent of the presence of H3K9me2 or DNA methylation (Mathieu et al., 2005), and in wheat mutually exclusive with H3K9me2 (Liu et al., 2021), suggesting that H3K27me2 and RdDM are independent of each other. In line with this idea, in maize, high H3K27me2 levels are observed at both RdDM and non-RdDM loci (Gent et al., 2014). In maize heterochromatin, however, H3K27me2 largely overlaps with H3K9me2 and DNA methylation, except that H3K27me2 is found at more loci (Gent et al., 2014). Links between small-RNA mediated transcriptional silencing and H3K27 methylation have been observed in protozoa. In *Tetrahymena thermophila*, small-RNA dependent recruitment of H3K27me3 to developmentally transcriptionally silenced loci has been observed (Liu et al., 2007). A strong relation between H3K27me2 enrichment and small RNA-mediated transcriptional silencing has been observed in the protozoan *Entamoeba histolytica* (Foda and Singh, 2015). Here we show that in *B’ mop1-1* and *mop3-1* plants, the H3K27me2 levels at the hepta-repeat are significantly lower than in *B’* wild type plants, suggesting that in maize MOP factors can mediate RNA-directed histone methylation.

Our observation of *mop* mutants resulting in reduced H3K27me2 and H3K9me2 levels does not resolve the relationship and potential hierarchy between these two histone modifications. It is also possible that the MOP siRNA pathway recruits more than one histone methyltransferase. In this scenario, both H3K27 and H3K9 could be direct targets, whereby the respective histone methyltransferases may be recruited at different levels in different tissues. Regardless of which histone modifier could be part of the MOP siRNA pathway at the *b1* hepta-repeat, it is likely not an anomaly. Our finding that 2% of *mop1* siRNA loci have low levels of mCHH is consistent with a common function of 24-nt siRNAs in something other than directing DNA methylation.

### Reinforcing cycle between mCHG and H3K9me2

How could a large role for siRNAs in directing histone modifications go unnoticed? For one reason, it is difficult to separate DNA methylation from histone modifications, especially given the massive pleiotropic effects of mutants. At the *b1* hepta-repeat, however, we observed a tissue-independent decrease in H3K9me2 in *mop3-1*, while mCHG remains high. mCHG is normally tightly linked to H3K9me2 because chromomethylases (CMTs) bind H3K9me2 containing nucleosomes and methylate cytosines in the CHG context (Bartee et al., 2001; Du et al., 2014, 2012; Lindroth et al., 2001; Stroud et al., 2014, 2013). In addition, H3K9 histone methyltransferases bind to methylated cytosines to methylate H3K9 (Du et al., 2014; Johnson et al., 2007). In maize, the CMTs ZMET2 and ZMET5 are required for mCHG (Fu et al., 2018; Gent et al., 2014; Li et al., 2014; Papa et al., 2001); ZMET2 also binds H3K9me2 (Du et al 2012). At the *B’* hepta-repeat, CHG methylation is, however, not decreased in *mop1-1* and *mop3-1* mutants, while H3K9me2 is significantly decreased, suggesting that CHG methylation is not sufficient to recruit H3K9 methyltransferases to maintain high H3K9me2 levels at the *B’* hepta-repeat. This supports the idea that the MOP1 RdRP and especially MOP3 NRPD1 play a role in recruiting H3K9 methyltransferases. It is unclear, however, how the high mCHG levels are maintained. Possibly, the reduced levels of H3K9me2 are still sufficient for ZMET2/5 to maintain high mCHG levels.

### Gene activation in presence of repressive marks

Previously, we postulated that low DNA methylation levels and lack of H3K9me2 and H3K27me2 at the *B-I* hepta-repeat would allow the binding of transcription factors driving activation of the hepta-repeat enhancer (Haring et al., 2010; Louwers et al., 2009a). We showed that activation of the *B-I* hepta-repeat enhancer was associated with histone acetylation and chromosomal interactions between the hepta-repeat, TSS and additional regulatory sequences at the *b1* locus, ultimately resulting in tissue-specific transcriptional enhancement of *b1*. The repressive marks at the *B’* hepta-repeat, DNA methylation, H3K9me2, and H3K27me2 were hypothesized to hamper enhancer activation and block the formation of a multi-loop structure, resulting in low *b1* expression. Accordingly, the presence of these repressive epigenetic marks are also negatively correlated with enhancer activity in other systems (Ferrari et al., 2014; Plank and Dean, 2014; Schmitz et al., 2022; Sekhon et al., 2012; Xie et al., 2013). Here we show the remarkable finding that, in *mop1-1* and *mop3-1* mutant plants, the regulatory sequences at the *b1* locus can be activated in the presence of high levels of DNA methylation and nucleosomes. This activation is illustrated by increased *b1* RNA and H3ac levels, and the formation of a multi-loop structure at the *B’* epiallele (Figs. 1-3). The restoration of the *b1* expression is, however, only to about 15% (*mop1-1*) or 23% (*mop3-1*) of the *B-I* level (Figure 1). Accordingly, the H3ac levels are (slightly) lower than observed for *B-I*. We hypothesize that in *B’ mop1-1* or *mop3-1* husk tissue, a reduction in H3K9me2 and H3K27me2 levels at each hepta-repeat is sufficient to allow transcription factors and other protein factors involved in enhancer function to bind the hepta-repeat, albeit with a relatively low affinity. This then results in an intermediately active enhancer that can physically interact with the additional regulatory sequences, ∼107, ∼47 and ∼15 kb upstream of the TSS; together they mediate the formation of an active transcription complex at the *b1* promoter, upregulating *b1* transcription. Alternatively, there could be near 100% reduction of H3K9me2 and H3K27me2 at a limited number of *b1* alleles, activating *b1* expression in only a fraction of cells. In either case, the high DNA methylation levels at the hepta-repeat enhancer allow the binding of proteins involved in enhancer activation to some extent.

Unlike in the *mop1-1* and *mop3-1* mutants, in both the dominant and recessive *mop2* mutants there is no significant decrease in H3K9me2 and H3K27me2 at the hepta-repeat enhancer, and the DNA methylation levels are, if anything, even slightly increased. Retention of these repressive marks in *mop2* mutants correlates with a lack of enhancer activation, which is associated with the formation of a single loop structure and no significantly increased *B’* transcript levels in these mutants. *Mop2/Rmr7* codes for NRP(D/E)2a, the second largest subunit of Pol IV and Pol V (Sidorenko et al., 2009; Stonaker et al., 2009). NRP(D/E)2a has two paralogs, NRP(D/E)2b and NRPE2c. NRP(D/E)2b is also part of Pol IV and V while NRPE2c is only part of Pol V (Haag et al., 2014; Sidorenko et al., 2009). Nevertheless, neither of these paralogs is able to substitute for NRP(D/E)2a with respect to paramutation in terms of the establishment of DNA and histone methylation at the *b1* hepta-repeat. However, one of the paralogs may substitute for NRP(D/E)2a to maintain these marks. Arabidopsis contains one functional homologue of NRP(D/E)2a, (NRP(D/E)2; (Herr et al., 2005; Kanno et al., 2005; Onodera et al., 2005; Pontier et al., 2005)). DNA blot analysis on Arabidopsis mutants of NRP(D/E)2 indicated a slight decrease in DNA methylation at 5S genes. So far, no effects on histone modifications are reported. In conclusion, our results indicate that repression of the *B’* epiallele can be maintained independent of MOP2 but does require MOP1 and MOP3.

### Effect on chromatin structure by mop mutants

The different *mop* mutants all hamper paramutation and block RdDM, but affect the chromatin structure differently. In the *mop1-1* and *mop3-1* mutants, repression at the *B’* hepta-repeat enhancer is released to different extents, but not in *mop2* mutants. We know little about the effect of these mutants on the binding of the RdDM components that are still functional. We expect, like observed for Arabidopsis, the MOP1 RdRP to stably bind the largest subunit of Pol IV (NRPD1/Mop3/Rmr6) (Law et al., 2011; Mishra et al., 2021). The effect of a lack of a functional MOP1 on the binding of the remaining RdDM components to chromatin in a *mop1-1* mutant is unknown.

We hypothesize that in a *mop3* mutant, Pol V, but not Pol IV, is still functional, resulting in transcription of the *B’* hepta-repeat by Pol V in the absence of RdDM. This transcription may lead to the observed decrease of H3K9me2 and H3K27me2 at the *B’* hepta-repeat, allowing enhancer activation. In line with this hypothesis, in Arabidopsis the production of Pol V transcripts is independent of mutations affecting components in siRNA biogenesis, and *de novo* and maintenance DNA methylation (DICER1-4, RDR2, DRM2, MET1 and DDM1) (Wierzbicki et al., 2008).

As discussed above, we cannot exclude that in *mop2* mutants, NRP(D/E)2a is replaced by NRP(D/E)2b or 2c, which may still be able to repress the *B’* hepta-repeat. Alternatively, both Pol IV and Pol V become non-functional and in absence of their transcription activity, the *B’* hepta-repeat stays repressed. In Arabidopsis, which has only one NRP(D/E), it has been seen that in the absence of NRP(D/E)2, the largest subunit of Pol V is destabilized, while NRPD1 is not (Pontier et al., 2005). If also true in maize, the Pol V protein complex would no longer bind, while the Pol IV complex may be dysfunctional.

The dissimilar effect of *mop1*, *mop2* and *mop3* mutations on the repressed state of the *B’* hepta-repeat is in line with findings for *pl1*. In 2-10% of wild-type plants, *Pl’* spontaneously reverts to the *Pl-Rh* state (Hollick et al., 1995) and is therefore less stable than the *B’* state, for which no reversion to *B-I* has been observed. In *rmr6-1* (*mop3*) mutants, *Pl’* changes to *Pl-Rh* with an increased reversion rate of 36% (Hollick et al., 2005), and also in *mop1-1* mutants a higher reversion rate to *Pl-Rh* was observed than in wild-type plants (Dorweiler et al., 2000). In *rmr7* (*mop2*) mutants, however, *Pl’* does not revert to *Pl-Rh*, indicating that the *Pl’* epiallele is more stably repressed in an *nrp(d/e)2a* mutant than in wild-type plants (Stonaker et al., 2009).

### Epigenetic memory

For *mop1-1* and *rmr6* (*mop3*) mutant plants, in which the production of siRNAs is severely decreased (Arteaga-Vazquez et al., 2010; Erhard et al., 2009; Nobuta et al., 2008; Sidorenko et al., 2009; Stonaker et al., 2009), it has been shown that the epigenetic memory of the paramutagenic *B’* state is maintained (Dorweiler et al., 2000; Hollick et al., 2005). Upon crossing out the *mop* mutations, *B’* can again paramutate *B-I* and is again lowly expressed, indicating the RdDM pathway is not involved in maintaining the epigenetic memory. We hypothesize that mCG and mCHG play a crucial role in maintaining the *B’* epigenetic state through meiosis, as in all *mop* mutants, the *B’* hepta-repeat remains associated with high mCG and mCHG levels. DNA methylation has been shown to play a central role in the stable transgenerational inheritance of silent epigenetic states in plants (Fitz-James and Cavalli, 2022; Mathieu et al., 2007). In line with this, Calarco et al. (2012) showed that mCG and mCHG are largely maintained in the germline. We, however, cannot entirely exclude a role for H3K9me2 and H3K27me2, even when the *B’* hepta-repeat H3K27me2 levels are significantly reduced tissue-independently in *mop1-1* and *mop3-1* mutants, and H3K9me2 levels in the *mop3-1* mutant. H3K9 methylation has been indicated to play a role in transgenerational inheritance in *S. pombe*, *D. melanogaster* and *C. elegans* (Fitz-James and Cavalli, 2022), and immunocytological data suggests that H3K9me2 may be retained during meiosis (Underwood et al., 2018). Currently, there is no role reported for H3K27me2 in transgenerational inheritance in plants.

### Concluding remarks

This study showed that, in the presence of high DNA methylation, a reduction in the repressive H3K9me2 and H3K27me2 marks at the regulatory *b1* hepta-repeat is sufficient to allow partial activation of the hepta-repeat enhancer function. This indicates that the mere presence of DNA methylation is not sufficient to entirely block enhancer function and that high levels of H3K9me2 and H3K27me2 are involved in blocking enhancer function. We also showed that high mCHG levels can be maintained in the presence of significantly reduced H3K9me2 levels, suggesting the reduced H3K9me2 levels are still sufficient for ZMET2/5 to maintain high mCHG levels. Intriguingly, we show that in maize, the RdRP and largest subunit of RNA Pol IV involved in RdDM also play a role in the maintenance of H3K9me2 and H3K27me2. More experiments are necessary to show if this maintenance role is confined to a limited number of sequences or occurs genome-wide. Our results also emphasize the possibility of RNA-directed histone modification in plants, as occurs in diverse fungi and animals. Future studies will be needed to examine the role of the MOP siRNA pathway in H3K9me2 and H3K27me2, and if RNA-directed histone modifications play a role in paramutation.

## Methods

### Genetic Stocks and Plant Materials

The plants stocks used in this study (*B’/B’ Mop1/Mop1* (K55), *B-I/B-I Mop1/Mop1* (W23), *B’/B’ Mop1/mop1-1* (K55/W23), *Mop2 B’/mop2-1 B’* (W23/K55), *Mop2 B’/mop2-2 B’* (W23/K55), *B’/B’ Mop3/mop3-1* (W23/K55) were obtained from V.L. Chandler (University of Arizona, Tucson, AZ) and grown in greenhouse conditions. Homozygous *mop* mutants were obtained by crossing *B’/B’ mop1-1/Mop1* with *B’/B’ mop1-1/mop1-1, Mop2 B’/mop2-1 B’* with *mop2-1 B’/mop2-1 B’, Mop2 B’/mop2-2 B’* with *mop2-2 B’/mop2-2 B’,* and *B’/B’ Mop3/mop3-1* with *B’/B’ mop3-1/mop3-1*, respectively. Homozygous progeny were identified using PCR genotyping (*mop1-1*) followed by sanger sequencing (*mop2*) and/or scoring pigment levels (*mop1-1*, *mop3-1*) (Sidorenko et al., 2009; Sloan et al., 2014). Heterozygous *B’ Mop2/mop2-1* mutants for experiments were mainly obtained from a cross of *Mop2 B’/mop2-1 B’* with *mop2-1 B’/mop2-1 B’.* In addition, for a few RNA and DNA isolations also samples from a cross between wild type *B’* and homozygous *mop2-1 B’* plants were used (data in Supplemental Figure S1, S6; derivation is indicated in the figures). The seedling tissue used for ChIP experiments comprised of 1-month-old seedlings with the roots and exposed leaf blades removed. The husk tissues employed for RNA, ChIP and 3C consisted of all leaves surrounding the maize ear, whereby the tough, outer leaves were discarded (Haring et al., 2010; Louwers et al., 2009a). Husks were harvested at the time of silk emergence. For DNA methylation analysis, mainly leaf blade tissue served as input material.

### RNA Analysis

RNA blot analysis was performed as described previously (Louwers et al., 2009a; Haring et al., 2010). RNA was isolated from husk tissue and 10 μg of RNA was size-fractionated by formaldehyde gel electrophoresis, blotted and hybridized with probes against the *b1* or *Sam* gene (Louwers et al., 2009a). RNA blots were hybridized with probes recognizing the coding region of *b1* or *Sam*. Band intensities were quantified using a Storm 840 phosphorimager and ImageQuant software (GE LifeSciences) and relative transcript levels of *b1* were calculated by normalization to transcript levels of the *Sam* housekeeping gene.

### Chromatin Immunoprecipitation

ChIP-qPCR experiments were performed on husk and seedling tissue as previously described (Haring et al., 2010, 2007). Antibodies were used against H3K9ac/K14ac (Upstate #06-599), H3K9me2 (Cell signaling #4658), H3K27me2 (Cell signaling #9728) and histone H3 (Abcam #1791). For the no-antibody reactions, 20 μl Rabbit serum/ChIP sample (Sigma R9133) was used. The specificity of the H3K9me2 and H3K27me2 antibodies was validated by slot blots containing different dilutions of the target peptides (Egelhofer et al., 2011). All ChIP-qPCR experiments were done using PLAT SYBR QPCR 500 (Invitrogen/Fisher, # 11733046) on an Applied Biosystems™ 7500 Real-Time PCR System, repeated 2-4 times with chromatin derived from different plants (see legends Figures 2,6 and 7, and Supplemental Figures S3, S11, S11). Primers that were used for the qPCR analysis are listed in Supplemental Table S1. *Actin* and *copia* served to normalize ChIP-qPCR data.

To test if *B’*, *B-I*, *B’ mop1-1, B’ Mop2/mop2-1*, *B’ mop2-1*, *B’ mop2-2* and *B’ mop3-1* behaved similar for the enrichment of the measured histone modifications in either seedling or husk tissue, two-way Analysis of variance (ANOVA) tests were used. Given the limited sample number, we assumed all datasets to be normally distributed and having similar variances. When the ANOVA tests indicated significant differences between the different (epi)genotypes, a pairwise t-test with a Bonferroni correction for multiple testing was performed to uncover which individual pairs differ significantly. The output from the ANOVAs and pairwise t-tests is summarized in Supplemental Table S2.

The *Copia* sequence behaves, on average, as a non-RdDM locus (Parent et al., 2021) and could therefore be used to normalize ChIP-qPCR data obtained for *B’ mop1-1* plants. In previous experiments, *Copia* sequences were used to normalize ChIP-qPCR data obtained with antibodies recognizing H3K9me2 and H3K27me2 (Haring et al., 2010, 2007). The majority of *Copia* elements are depleted of siRNAs (Gent et al., 2013), indicating they are rarely targeted by RdDM. To validate that the specific type of *Copia* sequence used in our previous experiments was not targeted by RdDM and could be employed for normalization in *mop* mutant backgrounds, the 107-nt sequence was blasted to the maize B73 genome (version 3) using parameters optimized for low similarity (e-value 1e-15) and including low complexity regions. Of the 163 matches, the 97 that were greater or equal to 95-nt in length, carried at least part of both primer binding sites, and had sequence coverage in a bisulfite sequencing experiment (Gent et al., 2014) were selected. These 97 loci were examined for the presence of CHH methylation, a hallmark of RdDM. Their average CHH methylation level was 1.66%, which is very close to the 1.34% of CHH methylation at non-RdDM loci and significantly different from the average CHH methylation level at RdDM loci (14.51%). In addition, analysis of H3K9me2 data from the same study revealed that these loci had an H3K9me2 enrichment value of 1.10, which is characteristic of non-RdDM loci (H3K9me2 enrichment of 1.12) rather than RdDM loci (H3K9me2 enrichment of 0.55) (Gent et al., 2014). This indicated that our *Copia* control sequence behaves, on average, as a non-RdDM locus and that it could be used to normalize our ChIP-qPCR data obtained from wild-type and *mop* mutant plants.

### Chromatin Conformation analysis (3C)

3C analyses were performed on *B’ mop1-1* husk tissue as described previously (Louwers et al., 2009a, 2009b). The data for *B’ Mop1/Mop1* and *B-I Mop1/Mop1* have been published before (Louwers et al., 2009a) and are shown for comparison. The experiments on *B’ mop1-1* tissues were conducted at the same time and using the same methods as the experiments on *B’ Mop1/Mop1* and *B-I Mop1/Mop1* tissues. The primers and TaqMan probes used are listed in Louwers *et al*. (2009a) In 3C experiments, to correct for primer amplification efficiencies, for each primer pair the qPCR data was normalized to a random ligation control sample produced using a BAC clone containing the *b1* locus and an amplified control template for the *Sam* locus. To control for quantity and quality of the 3C samples, the data were normalized to 3C values measured for the *Sam* locus.

### DNA Methylation Analysis by DNA Blotting

DNA blot analysis was performed as described previously (Haring et al., 2010). Most DNA samples were derived from leaves collected at different stages of development (Supplemental Figures S4, S6 and S7). DNA was isolated (Dellaporta et al., 1983) and 5 μg of DNA was digested with different methylation sensitive restriction enzymes in combination with *Eco*RI or *Bam*HI according to the manufacturer’s specifications. After size fractionation by electrophoresis the gels were blotted onto membrane (Blotting-Nylon 66 membranes, Sigma-Aldrich # 15356), followed by UV fixation. The resulting blots were hybridized to a 853-nt repeat probe. The relative band intensities obtained for the enzymes *Hpa*I, *Pst*I, *Hha*I, *Hae*II *Alu*I, *Sau*96I, *Bsm*AI were quantified for all (*Hpa*I, *Hha*I, *Hae*II) or representative examples (*Pst*I, *Alu*I, *Sau*96I, *Bsm*AI), using a Storm 840 phosphorimager and ImageQuant software (GE LifeSciences). The most probable DNA methylation patterns were identified by the least squares fit comparing the relative band intensities with theoretical possible intensities derived from all possible options for DNA methylation patterns (Haring et al., 2010). *Bsm*AI generated relatively small fragments, including similarly sized 5’ and 3’ border fragments complicating the computational analysis. The degrees of DNA methylation indicated (Figure 4A, Supplemental Figure S6, S7) are our best estimates. To test for complete digestion, probes recognizing unmethylated DNA regions at the *b1* locus were used (probes A, D2 and 21, Figure 1 in Haring et al., 2010).

### DNA Methylation Analysis by Bisulfite Sequencing

For bisulfite sequencing, genomic maize DNA was extracted (Dellaporta et al., 1983) from leaf four of two *B’*, two *B-I* and three *B’ mop1-1* plants in a V4 stage (Ritchie et al., 1986), and 400 ng of DNA was treated with bisulfite using the EZ DNA Methylation-Gold kit (Zymo Research, D5006). The DNA regions of interest were PCR-amplified (10 min 95°C, followed by 40 PCR cycles (30 sec 95°C, 30 sec 50°C (repeat) or 52°C (*Fie2*), 30 sec 72°C), and 5 min at 72°C). The amplification was performed using MethylTaq DNA polymerase (Diagenode, C09010010), a forward (KL1310; TGGTGTTTAAAAATTYATGTTTTTGTG) and reverse primer (KL1844; TCCACRARTCATCRTCTCCAAACA) for a 320-nt repeat junction fragment and a forward (AAGATTTGAGATTYGATTTGAAGTGTG) and reverse primer (ACTTTCCCCTCCRCCTAATTCTCCTTA) for a 226-nt *Fie2* fragment that served as a control for complete bisulfite conversion (similar to the −302 to −91 fragment described in Gutierrez-Marcos et al., 2006). PCR fragments were cloned into the pJET 1.2 vector (CloneJet PCR Cloning Kit, Thermo Scientific) following the manufacturer’s instructions.

Clones carrying inserts of the correct size were identified by PCR. DNA was isolated from positive clones (GeneJet Plasmid Miniprep Kit, Thermo scientific) and subjected to Sanger sequencing. To determine the percentage, sequence context and pattern of cytosine methylation, sequences were analyzed using Kismeth (Gruntman et al., 2008). For each DNA sample approximately 15 clones were sequenced for *Fie2*, and 30 clones for the repeat junction region.

### Genomic Alignments and Data Analysis of DNA methylation-seq and Small RNA Reads

Small RNA sequences from developing ear were obtained from the Sequence Read Archive, runs SRR1583941 and SRR1583942 for the *mop1-1* mutant; and SRR1583943 and SRR1583944 for wild-type (Gent et al., 2014). Both were in a genetic background closely related to B73. The methylC-seq data from the same study were derived from pure B73 developing ear, runs SRR1583945, SRR1583946, and SRR1583947. The Enzymatic Methyl-seq (EM-seq) data from developing second leaf of B73 were from the NAM founders assembly project and available through the European Nucleotide Archive, runs ERR5347654, ERR5347655, and ERR5347656 (Hufford et al., 2021).

Both methylC-seq and EM-seq reads were processed and mapped with BS-Seeker2 (Guo et al., 2013) according to published methods (Hufford et al., 2021), except that no mismatches were allowed (−m 0) and mapping was done in end-to-end mode (−-bt2--end-to-end). Potential PCR duplicates were removed from the ear methylC-seq reads using the BS-Seeker2 FilterReads tool. Methylation values and coverage for genomic loci were obtained using the CGmapTools mtr tool for each of the three sequence contexts separately (Guo et al., 2018). For defining low mCHH and high mCHH *mop1* loci, a minimum effective coverage of 20 was required, where effective coverage is the sum of the read coverage of each individual cytosine in each locus. Methylation values were measured as the average of each site in each locus. Low mCHH *mop1* loci were defined as having less than 0.02 mCHH, and high mCHH *mop1* loci as having greater than or equal to 0.02 mCHH in the source B73 ear methylome (Gent et al., 2014). These were initially identified based on 50-bp non-overlapping intervals, but all adjacent loci that met criteria for low mCHH were merged, and all adjacent loci that met criteria for high mCHH were also merged. Only merged loci consisting of at least two adjacent loci were included in the final sets, giving them minimum lengths of 100 bp. The methylation of these loci was reevaluated using the leaf methylome (Hufford et al., 2021), but with a constraint that only loci with at least five cytosines of the specific cytosine context being spanned by reads were included in the analysis.

Small RNA-seq reads were quality filtered, trimmed of adapters, and filtered for lengths of 20-25 nt using cutadapt (Martin, 2011), parameters -q 20 -a GATCGGAAGAGCACACGTCT -e .05 −O 5 --discard-untrimmed -m 20 -M 25. Reads were aligned to the genome with Bowtie2, –very-sensitive parameters (Langmead and Salzberg, 2012). Reads that overlapped at least 90% of their lengths with tRNA, 5S RNA, NOR, or miRNA loci were removed using the BEDTools intersect tool with parameters -v -f .9 (Quinlan and Hall, 2010). These excluded loci were identified using published methods (Gent et al., 2022), but applied to the B73 v5 reference genome (Hufford et al., 2021). The read counts for miRNA loci were used for normalization in downstream steps. The two input files for wild-type and the two for the *mop1-1* mutant were combined into single files using the SAMtools merge tool (Li et al., 2009) Uniquely-mapping reads were selected using the SAMtools view tool with the -q20 parameter. Perfect-matching siRNAs were selected using grep for the pattern ‘NM:i:0’. 24-nt siRNAs were selected from the complete set of 20 to 25-nt siRNAs using grep for the pattern ‘24M’. Read coverage over 50-bp intervals of the genome was obtained using the BEDTools coverage tool. All intervals were identified that had at least 25 overlapping reads and which spanned at least 30 bp of the interval from wild-type. In addition, they had to have a minimum of ten-fold fewer overlapping reads in the *mop1-1* mutant than in wild-type (normalized by the number of miRNA reads). These, called *mop1* loci, were then further filtered using methylation data as to generate low mCHH and high mCHH *mop1* loci as described above.

## Supporting information

Hovel-Supplemental Figures

## Gene IDs

*mop1*: Zm00001eb080370

*mop2/rmr7*: Zm00001eb068960

*mop3/rmr6*: Zm00001eb035080

## Acknowledgements

We would like to thank Vicki Chandler and Lyudmila Sidorenko for providing seeds. We are grateful to Nicole Riddle and Sarah Gadel who tested the specificity of the H3K9me2 and H3K27me2 Cell signaling antibodies with Slot Blots. Shiv Grewal is thanked for advice and fruitful discussion, Ludek Tikovsky and Harold Lemereis for taking excellent care of the maize plants, and Christel Rooker for proofreading of the manuscript. I.H. was supported by the Systems Biology Research Priority fund of the University of Amsterdam. K.P. is supported by the Topsector Horticulture & Starting materials; J.I.G was supported by a National Science Foundation grant (2114797). M.S. was supported by the Royal Netherlands Academy of Arts and Sciences (KNAW).

## Author contributions

I.H., M.L., M.H., J.I.G. and M.S. designed the research; R.B., I.H., M.L., M.H and J.I.G. performed research, analyzed data and made figures; K.P. performed statistical analysis and generated figures; I.H., J.I.G. and M.S. wrote the manuscript.

## Supplemental data

**Supplemental Figure S1.** *b1* expression levels in maize husk tissues.

**Supplemental Figure S2.** Schematic representation of the *b1* and *Sam* locus.

**Supplemental Figure S3.** *B’* hepta-repeat is marked by H3 acetylation in *mop1-1* and *mop3-1* mutants.

**Supplemental Figure S4.** In *mop* mutants slight differences in DNA methylation were observed at the *B’* repeat junction regions by DNA blot analysis.

**Supplemental Figure S5.** Consensus double-stranded DNA sequence of the 2nd-6th repeat. **Supplemental Figure S6.** Detailed DNA methylation data for *B’* in *mop* mutants using methylation sensitive restriction enzymes and DNA blot analysis.

**Supplemental Figure S7.** DNA methylation levels at *B’* hepta-repeat do not progressively decrease during development of a *B’ mop1-1* plant.

**Supplemental Figure S8.** Targeted bisulfite data for *B’*, *B-I*, *B’ mop1-1*, *B’ Mop2/mop2-1* and *B’ mop3-1* DNA derived from leaf 4 of V4 stage plants.

**Supplemental Figure S9.** Identification of low and high mCHH *mop1* loci and their respective siRNA levels.

**Supplemental Figure S10.** No decrease in nucleosome occupancy was observed in the *mop* mutants.

**Supplemental Figure S11.** Transcriptional activation in *mop1-1* and *mop3-1* is associated with reduced levels of repressive histone marks.

**Supplemental Table S1.** Primers used in ChIP-qPCR experiments.

**Supplemental Table S2.** Statistical analyses of the ChIP-qPCR experiments.

---

**Table S1.**
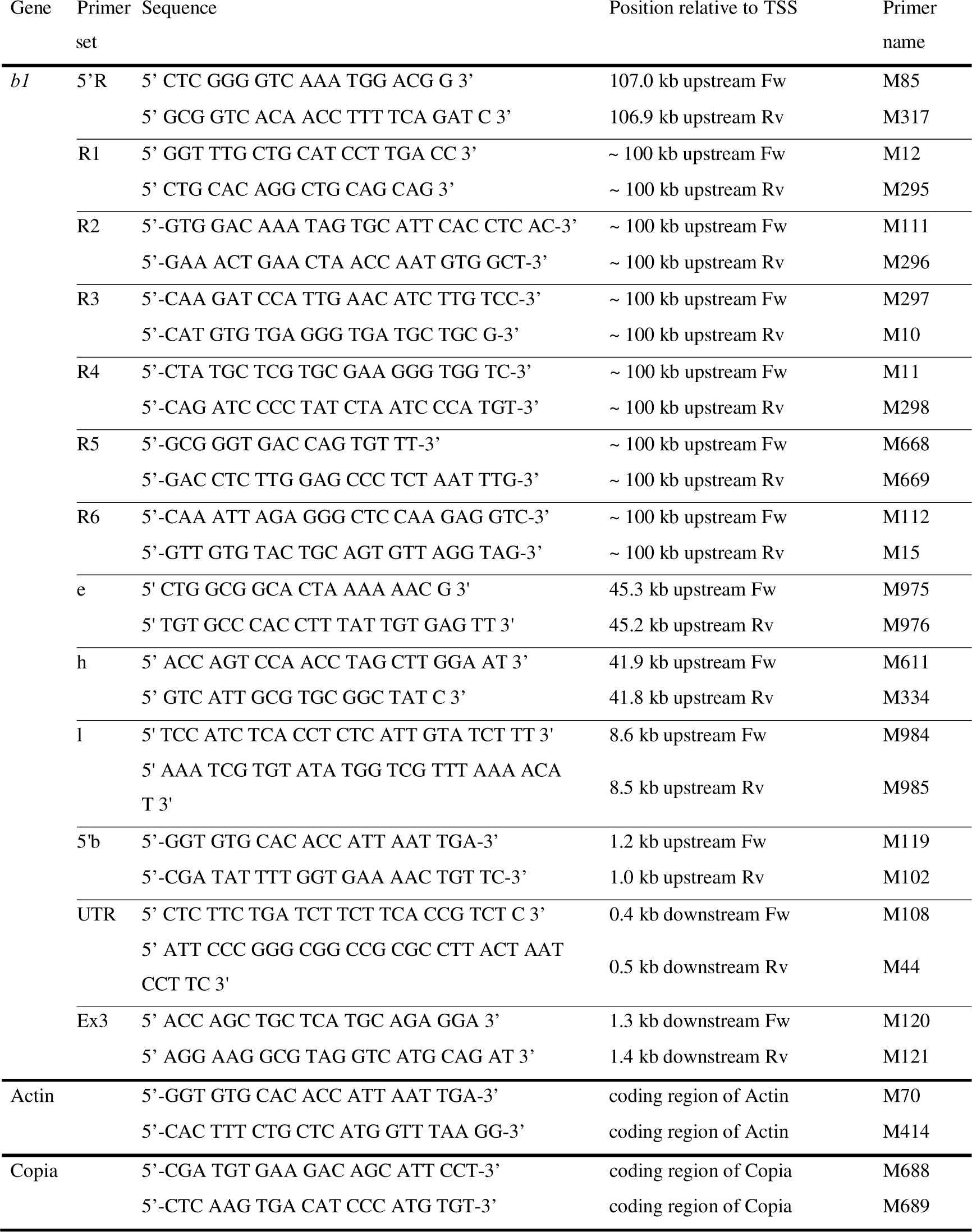
Primers used in ChIP-qPCR experiments.

## Notes

### Competing Interest Statement

The authors have declared no competing interest.

### Summary of Updates

This version has been revised to add the supplemental material

