## Supplementary material for "RdDM mutants reduce histone methylation rather than DNA methylation at the paramutated maize *b1* enhancer": Hovel-Supplemental Figures

### Supplemental Figure S1

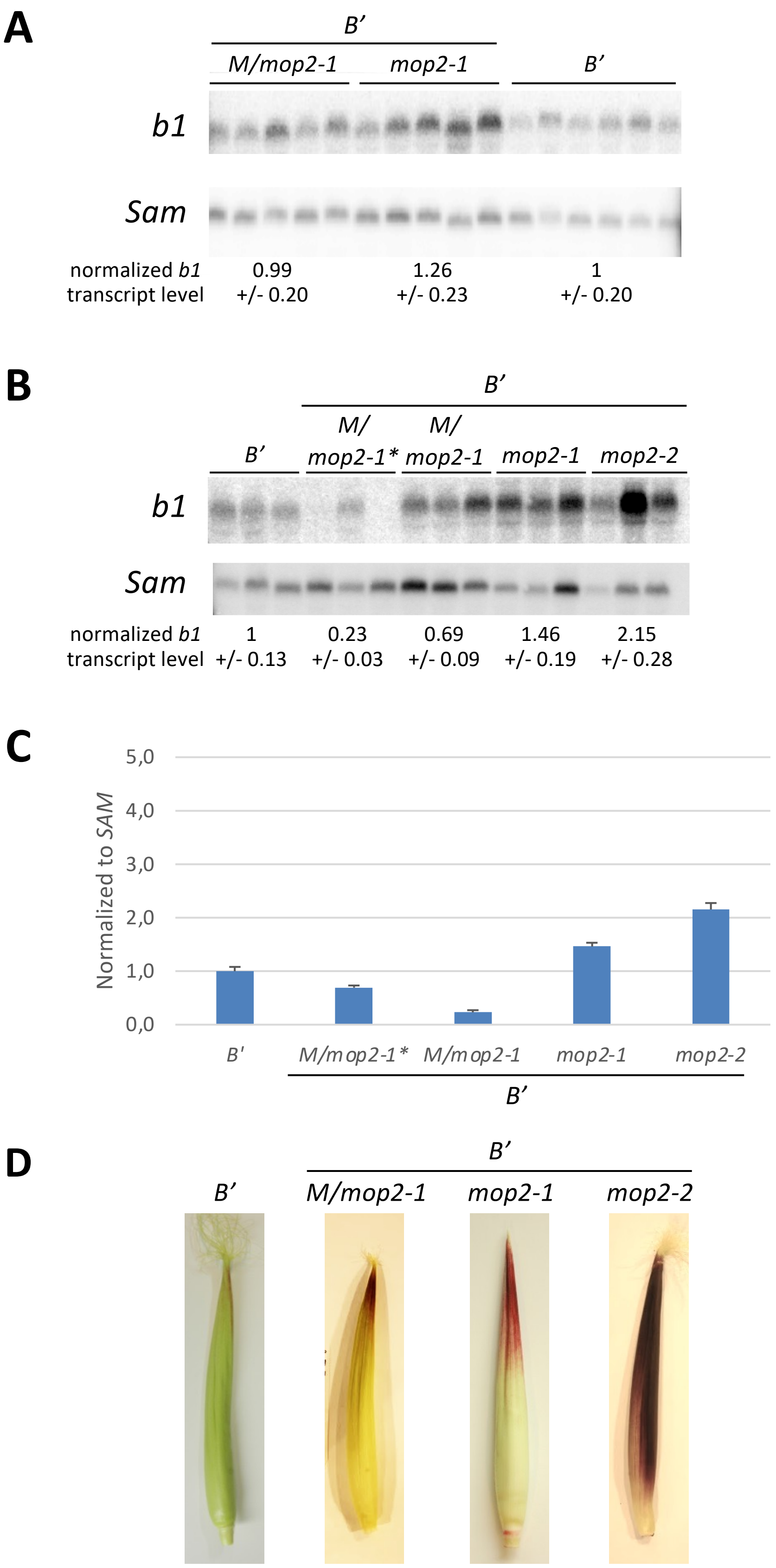

**Supplemental Figure S1.** *b1* expression levels in maize husk tissues. (A-C) RNA blot analysis of RNA from husk tissue of *B'*, *B'Mop2/mop2-1 (B'M/mop2-1), B'mop2-1, B'mop2-2,* and *B'Mop2/mop2-1* derived from a *B'* x *B'mop2-1* cross (*B'M/mop2-1**), using probes recognizing the coding region of *b1* (exons 7-9) and *Sam* (see Supplemental Figure S2). (A, B) RNA blots. The intensities of the bands representing full-length transcripts were quantified. The values below the blots represent the average plus standard deviation of *b1* transcript levels normalized to *Sam* transcript levels. (C) The quantification of the RNA blot shown in (B) shown in a graph.

### Supplemental Figure S2

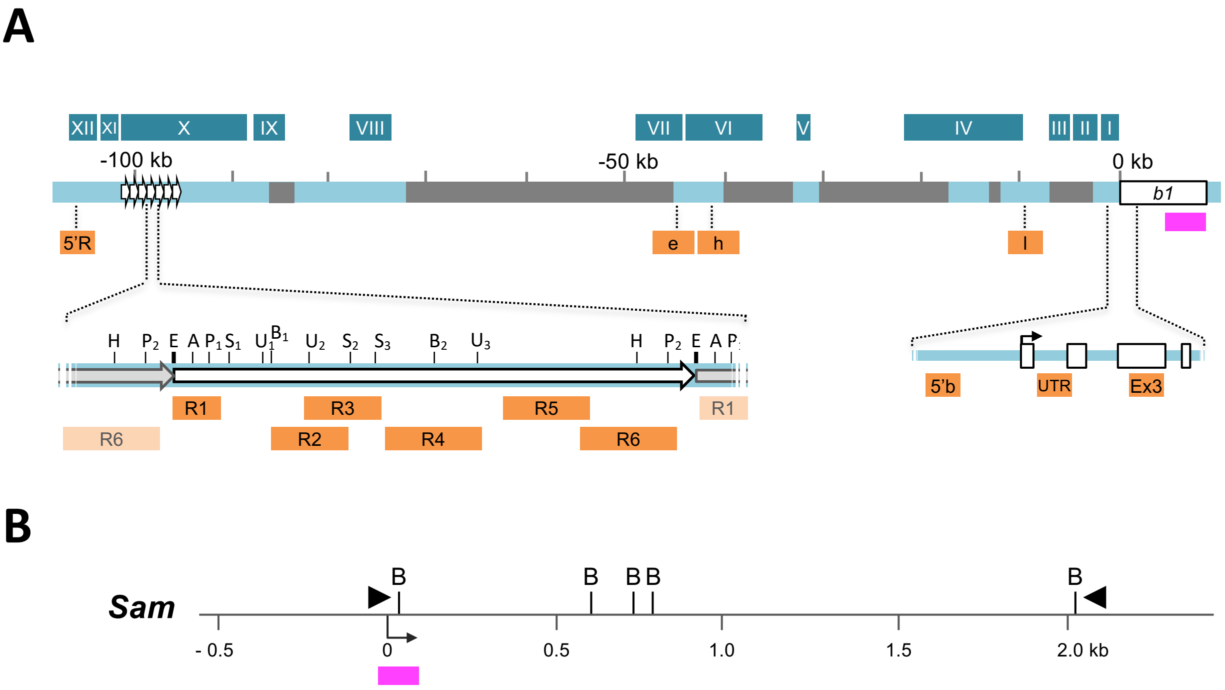

**Supplemental Figure S2.** Schematic representation of the *b1* and *Sam* locus. A) Schematic representation of the *b1* locus, indicating the regions and sites monitored in ChIP, 3C and DNA methylation analyses. The white box labeled *b1* depicts the *b1* coding region, empty white boxes in zoom below represent exons, the hooked arrow the TSS (0 kb); the *b1* 5'UTR consists of exon 1 and intron 1. Arrows 100 kb upstream represent the hepta-repeat, and dark grey boxes transposable elements. The *Bgl*II fragments analyzed in 3C experiments are indicated with dark blue boxes and white Roman numerals. The amplicons monitored in ChIP-qPCR experiments are indicated with orange boxes. Due to sequence divergence, primer sets R1 and R5 do not detect the 1st and 7th repeat, respectively. Restriction sites are indicated by letters: H, *Hpa*I; P, *Pst*I; E, *Hha*I and *Hae*II; A, *Alu*I; S, *Sau*96I; U, *Sau*3AI; B, *Bsm*AI; Numbers discriminate individual sites present more than once every repeat. Magenta box, probe for RNA analysis.
B) Schematic representation of the *Sam* locus, indicating the *Bgl*II fragments and amplicons analyzed in the 3C experiments, respectively. *Sam* is used for data normalization. B, *Bgl*II sites; triangles, 3C primers; hooked arrow, transcription start site; Magenta box, probe for RNA analysis (Figure 1 and Supplemental Figure S1).

### Supplemental Figure S3

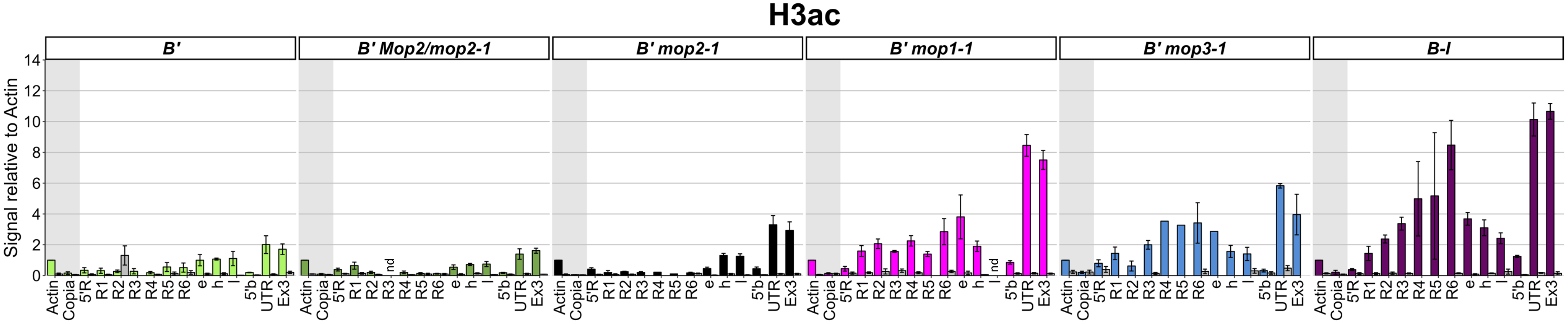

**Supplemental Figure S3.** *B'* hepta-repeat is marked by H3 acetylation in *mop1-1* and *mop3-1* mutants. ChIP experiments were performed on husk tissue from *B'* wild-type (light green), *B'* in a *mop1-1* (magenta), *Mop2/mop2-1* (dark green), *mop2-1* (black), and *mop3-1* (blue) background, and *B-I* plants (dark purple), using an anti-H3ac antibody and normalization to actin values. Colored bars indicate the ChIP signals, grey bars indicate signals obtained for the no-antibody immunoprecipitations. The error bars indicate the standard error of the mean (SEM) of three (*B', B' mop1-1, B' Mop2/mop2-1*, or four (*B' mop2-1*, *B' mop3-1, B-I)* replicate experiments. nd = no data for primer pair.

### Supplemental Figure S4

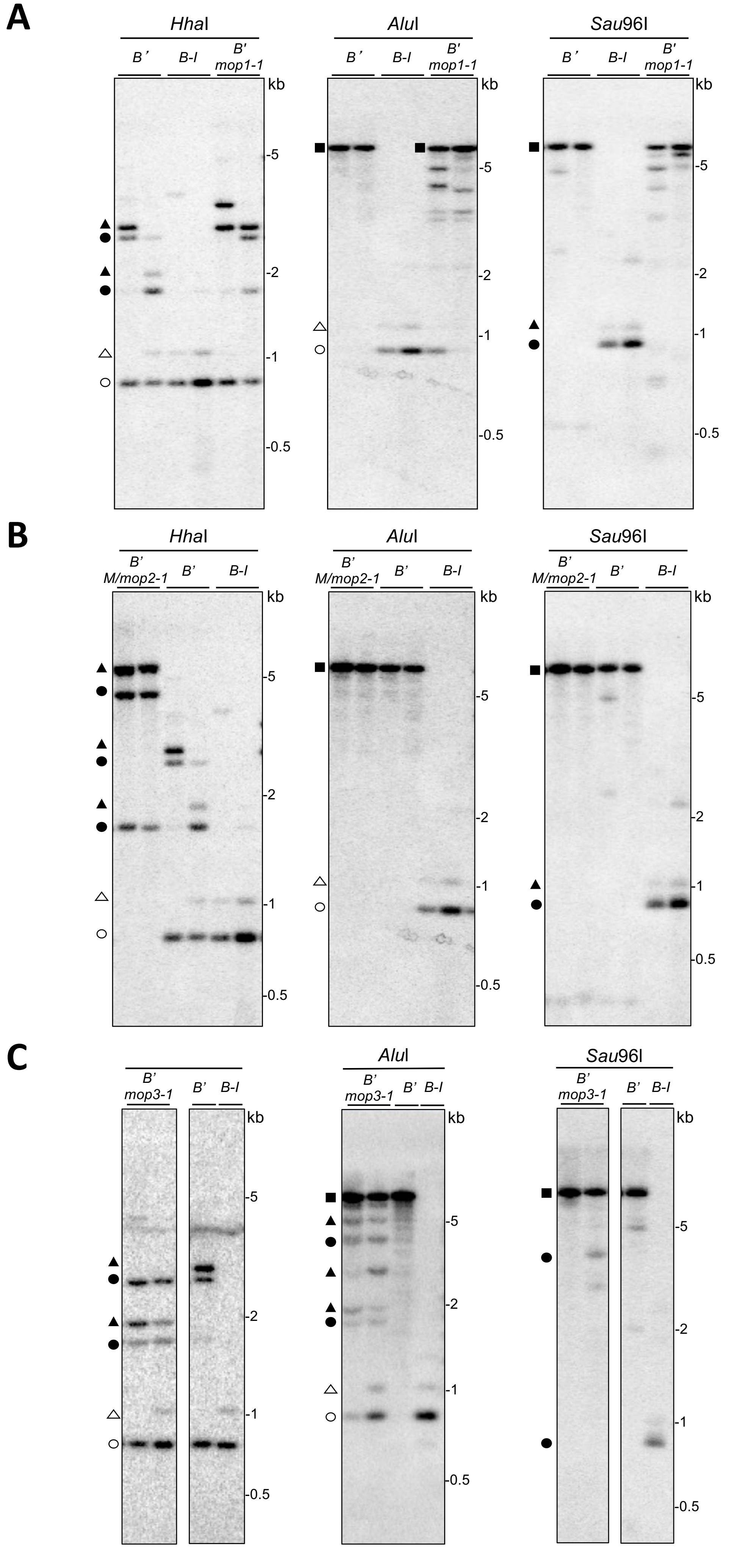

**Supplemental Figure S4.** In *mop* mutants slight differences in DNA methylation were observed at the *B'* repeat junction regions by DNA blot analysis. Genomic DNA was digested with *Eco*RI and the methylation-sensitive enzymes indicated. Representative examples of blots hybridized with the repeat probe are shown. Circles and triangles indicate fragments comprising repeat sequences and repeat plus 3' flanking sequences, respectively (Haring et al 2010). Open and filled symbols indicate fragments resulting from complete and incomplete digestion, respectively. Filled squares indicate fragments in which nearly all recognition sites are methylated. The *B'* and *B-I* patterns are shown as comparison. A) *B' mop1-1*, B) *B' Mop2/mop2-1 (M/mop2-1 ),* C) *B' mop3-1.*

### Supplemental Figure S5

```
                                                AluI          PstI
ccatgggtttgctgcatccttgacccgtagcctcactcacagctatgcaactgctgcagcctgtgcaggc 70
ggtacccaaacgacgtaggaactgggcatcggagtgagtgtcgatacgttgacgacgtcggacacgtccg

                    Sau96I
ttagcctcagcctatcgtggcccgacaactaaacaggtcgtgcagtgctctccccaagtcccgaccacta 140
aatcggagtcggatagcaccgggctgttgatttgtccagcacgtcacgagaggggttcagggctggtgat

          Sau3AI      BsmAI
atagtcgtgatccctgtttggagacgatgactcgtggacaaatagtgcattcacctcaccctcacacata 210
tatcagcactagggacaaacctctgctactgagcacctgtttatcacgtaagtggagtgggagtgtgtat

                    Sau3AI
ttttttttttgaaatcaagatccattgaacatcttgtccagttaaatcactggacaccgtgacagccaca 280
aaaaaaaaaactttagttctaggtaacttgtagaacaggtcaatttagtgacctgtggcactgtcggtgt

                       Sau96I                                  Sau96I
ttggttagttcagttcgtggtggaccgatggttcgcagtcgcagcatcaccctcacacatggtccgcatg 350
aaccaatcaagtcaagcaccacctggctaccaagcgtcagcgtcgtagtgggagtgtgtaccaggcgtac

gctacgcgtatctatgttcgtgcgaagggtggttctatatatttagactaccttcagtggatagggtaaa 420
cgatgcgcatagatacaagcacgcttcccaccaagatatataaatctgatggaagtcacctatcccattt

                            BsmAI
attaataaaggacacataattatagatgagactatttaatattttttttttacataaatgagatttgaaat 490
Taattatttcctgtgtattaatatctactctgataaattataaaaaaaaatgtatttactctaaacttta

                             Sau3AI
gatgactatgacatgggattagataggggatctggattattgcagggtgttaaatcctgagcgattttcg 560
ctactgatactgtaccctaatctatcccctagacctaataacgtcccacaatttaggactcgctaaaagc

cgggtgaccagtgttttttttaccgttttcgatactagacgaaatcaaatttcaaaaatctcacttaaaa 630
gcccactggtcacaaaaaaaatggcaaaagctatgatctgctttagtttaaagtttttagagtgaatttt

tttttattttaaaaaaaaaccataaaaatgggtaaaaatttttattacaaattagagggctccaagaggt 700
aaaaataaaatttttttttggtatttttacccatttttaaaaataatgtttaatctcccgaggttctcca

                                                                HpaI
ctataaaaatttggtgtttaAaaattcatgtttttgtgccaaaaaacaaaaaggttctagacctgttaac 770
gatatttttaaaccacaaattttaagtacaaaaacacggttttttgtttttccaagatctggacaattg

                                                  PstI
ccaaatggatattgttgcatctcccccaaattctttatctacctaacactgcagtacacaacttttatac 840
ggtttacctataacaacgtagagggggtttaagaaatagatggattgtgacgtcatgtgttgaaaatatg

     HhaI/HaeII
cgaatacttggcgcca 856
gcttatgaaccgcggt
```

**Supplemental Figure S5.** Consensus double-stranded DNA sequence of the 2nd-6th repeat. Red indicates cytosines in CG context, blue in CHGs, green in CHH context. The last three nucleotides (cca) shown are the first three nucleotides of the next repeat.

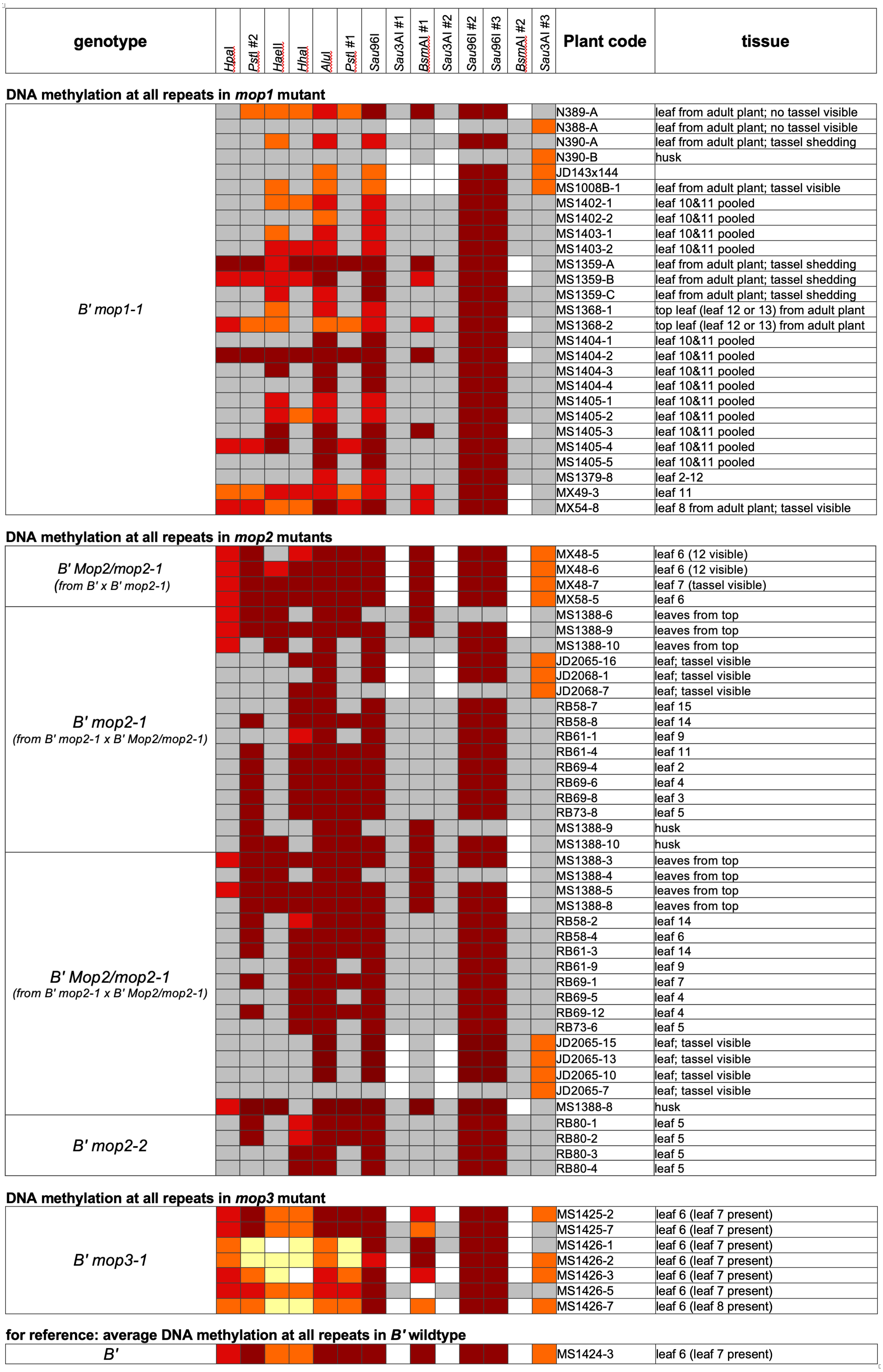

**Supplemental Figure S6.** Detailed DNA methylation data for *B'* in *mop* mutants using methylation sensitive restriction enzymes and DNA blot analysis. Every site checked for a specific sample is indicated. Grey cells indicate restriction sites not measured for particular samples. The degree of DNA methylation at the various sites is indicated by color-coding: 0-12.5%, white; 37.5-62.5%, orange; 62.5-87.5%, red; 87.5-100% methylation, dark red. The *B'* consensus pattern (Haring et al, 2010) is indicated at the bottom. See Fig. 4A for location of the sites. For details on the quantification of the DNA methylation levels see Haring *et al.* (2010).

### Supplemental Figure S7

A

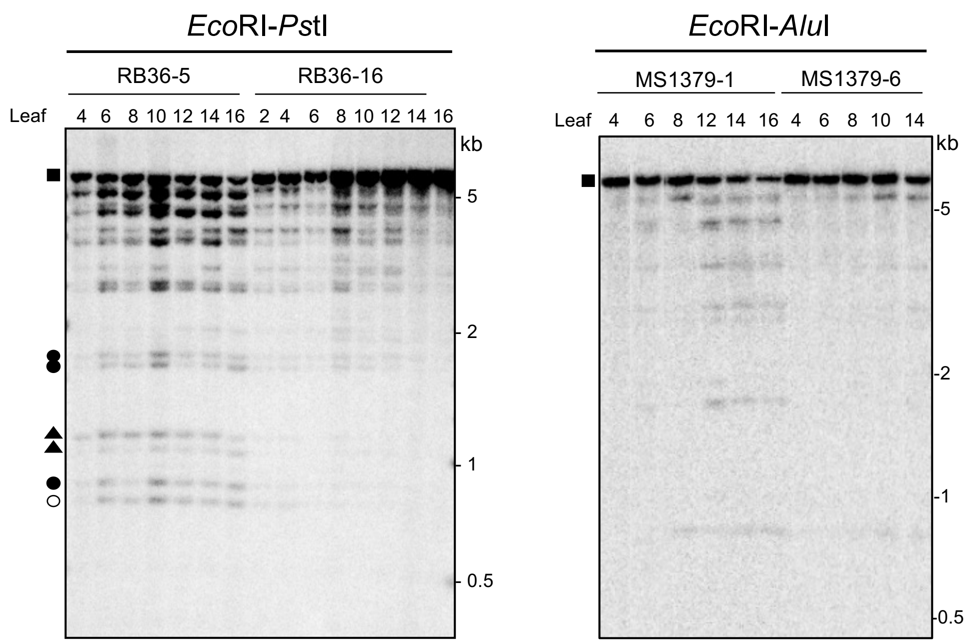

B

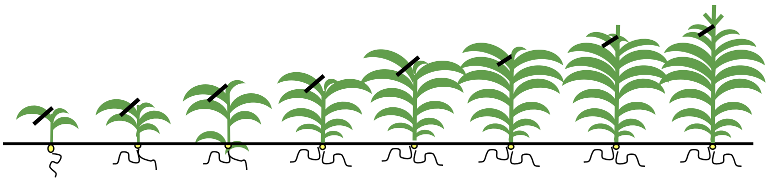

| genotype | Plant | Enzyme | Leaf 2 | Leaf 4 | Leaf 6 | Leaf 8 | Leaf 10 | Leaf 12 | Leaf 14 | Leaf 16 |
|---|---|---|---|---|---|---|---|---|---|---|
| ***B'*** | | *Sau*96I | | | | | | | | |
| *B' mop1-1* | MS1379-7 | *Sau*96I | | | | | | | | NA |
| *B' mop1-1* | MS1379-8 | *Sau*96I | | | | | | | | NA |
| ***B'*** | | *Alu*I | | | | | | | | |
| *B' mop1-1* | MS1379-1 | *Alu*I | NA | | | | NA | | | |
| *B' mop1-1* | MS1379-6 | *Alu*I | NA | | | | | | | NA |
| *B' mop1-1* | MS1379-8 | *Alu*I | | | | | | | | NA |
| *B' mop1-1* | RB36-5 | *Alu*I | NA | NA | | | | | | |
| *B' mop1-1* | RB36-16 | *Alu*I | NA | | | | | | | |
| ***B'*** | | *Pst*I | | | | | | | | |
| *B' mop1-1* | MS1379-1 | *Pst*I | | | | | | | | |
| *B' mop1-1* | MS1379-6 | *Pst*I | NA | | | | | | | NA |
| *B' mop1-1* | RB36-5 | *Pst*I | NA | | | | | | | |
| *B' mop1-1* | RB36-16 | *Pst*I | | | | | | | | |

**Supplemental Figure S7.** DNA methylation levels at *B'* hepta-repeat do not progressively decrease during development of a *B' mop1-1* plant. A) Genomic DNA was digested with *Eco*RI and the methylation-sensitive enzymes indicated. Representative examples of blots hybridized with the repeat probe are shown. Circles and triangles indicate fragments comprising repeat sequences, and repeat plus 3' flanking sequences, respectively (Haring *et al.* 2010). Open and filled symbols indicate fragments resulting from complete and incomplete digestion, respectively. Filled squares indicate fragments in which nearly all recognition sites are methylated. B) Detailed representation of the DNA methylation data obtained for leaves collected during development of *B' mop1-1* and as a comparison *B'* plants. The degree of DNA methylation is indicated by color-coding: 37.5-62.5%, orange; 62.5-87.5%, red; 87.5-100% methylation, dark red; NA, not analyzed. The specific plants of the families analyzed (MS1379 and RB36) are indicated. Leaves were collected when the consecutive leaf had appeared. For details on the quantification of the DNA methylation levels see Haring *et al.* (2010).

A

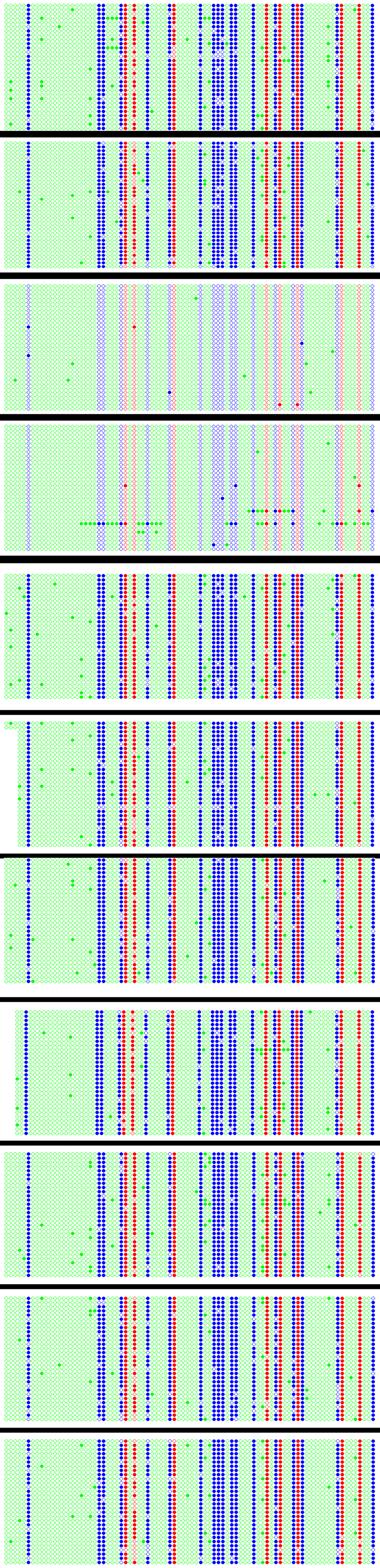

| *repeat* | single clones of bisulfite converted DNA | total number of cytosines | methylated fraction |
|---|---|---|---|
| *B'* sample 1 | | CG (240) | **94.58%** |
| | | CHG (540) | **92.22%** |
| | | CHH (1740) | **2.75%** |
| | | All (2520) | **30.67%** |
| *B'* sample 2 | | CG (272) | **91.54%** |
| | | CHG (612) | **89.37%** |
| | | CHH (1972) | **1.77%** |
| | | All (2856) | **29.10%** |
| *B-I* sample 1 | | CG (248) | **1.20%** |
| | | CHG (588) | **0.71%** |
| | | CHH (1794) | **0.44%** |
| | | All (2600) | **0.58%** |
| *B-I* sample 2 | | CG (240) | **3.33%** |
| | | CHG (539) | **2.96%** |
| | | CHH (1740) | **1.95%** |
| | | All (2519) | **2.30%** |
| *B' mop1-1* sample 1 | | CG (240) | **97.08%** |
| | | CHG (540) | **92.96%** |
| | | CHH (1736) | **2.24%** |
| | | All (2516) | **30.76%** |
| *B' mop1-1* sample 2 | | CG (240) | **93.75%** |
| | | CHG (539) | **93.32%** |
| | | CHH (1655) | **1.69%** |
| | | All (2434) | **31.06%** |
| *B' mop1-1* sample 3 | | CG (240) | **96.25%** |
| | | CHG (539) | **92.2%** |
| | | CHH (1736) | **1.78%** |
| | | All (2515) | **30.18%** |
| *B' Mop2/mop2-1* sample 1 | | CG (240) | **94.58%** |
| | | CHG (540) | **93.88%** |
| | | CHH (1650) | **2%** |
| | | All (2430) | **31.56%** |
| *B' Mop2/mop2-1* sample 2 | | CG (240) | **96.25%** |
| | | CHG (540) | **93.88%** |
| | | CHH (1739) | **2.70%** |
| | | All (2519) | **31.16%** |
| *B' mop3-1* sample 1 | | CG (240) | **94.58%** |
| | | CHG (540) | **89.44%** |
| | | CHH (1739) | **1.49%** |
| | | All (2519) | **29.22%** |
| *B' mop3-1* sample 2 | | CG (240) | **91.25%** |
| | | CHG (540) | **93.14%** |
| | | CHH (1739) | **1.49%** |
| | | All (2519) | **29.70%** |

●○ CG ●○ CHG ●○ CHH

B

| *Fie2* | single clones of bisulfite converted DNA | total number of Cs | non-converted fraction |
|---|---|---|---|
| *B'* sample 1 | 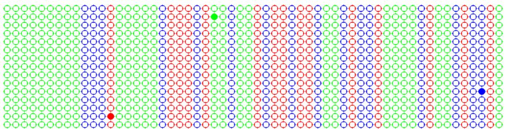 | CG (224) | **0.44%** |
| | | CHG (240) | **0.41%** |
| | | CHH (395) | **0.25%** |
| | | All (859) | **0.35%** |
| *B'* sample 2 | 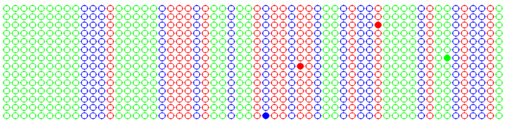 | CG (210) | **0.95%** |
| | | CHG (224) | **0.44%** |
| | | CHH (378) | **0.26%** |
| | | All (812) | **0.49%** |
| *B-I* sample 1 | 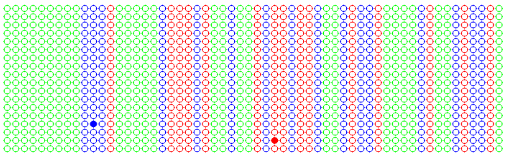 | CG (270) | **0.37%** |
| | | CHG (288) | **0.34%** |
| | | CHH (471) | **0%** |
| | | All (1029) | **0.19%** |
| *B-I* sample 2 | 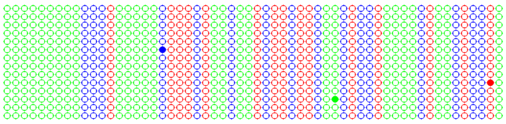 | CG (210) | **0.47%** |
| | | CHG (224) | **0.44%** |
| | | CHH (374) | **0.26%** |
| | | All (808) | **0.37%** |
| *B' mop1-1* sample 1 | 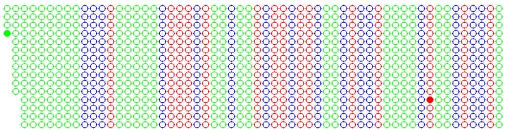 | CG (225) | **0.44%** |
| | | CHG (240) | **0%** |
| | | CHH (387) | **0.25%** |
| | | All (852) | **0.23%** |
| *B' mop1-1* sample 2 | 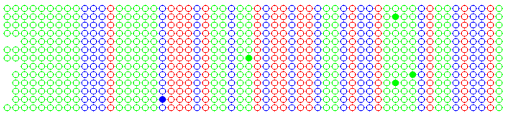 | CG (195) | **0%** |
| | | CHG (208) | **0.48%** |
| | | CHH (341) | **1.17%** |
| | | All (744) | **0.67%** |
| *B' mop1-1* sample 3 | 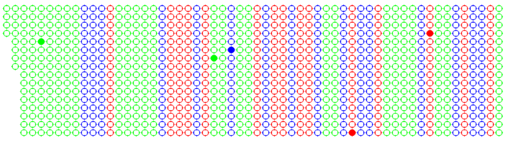 | CG (240) | **0.83%** |
| | | CHG (256) | **0.39%** |
| | | CHH (412) | **0.48%** |
| | | All (908) | **0.55%** |
| *B' Mop2/mop2-1* sample 1 | 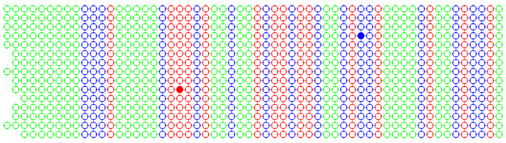 | CG (224) | **0.44%** |
| | | CHG (238) | **0.42%** |
| | | CHH (389) | **0%** |
| | | All (851) | **0.24%** |
| *B' Mop2/mop2-1* sample 2 | 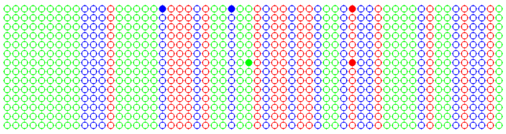 | CG (210) | **0.95%** |
| | | CHG (224) | **0.89%** |
| | | CHH (374) | **0.26%** |
| | | All (808) | **0.62%** |
| *B' mop3-1* sample 1 | 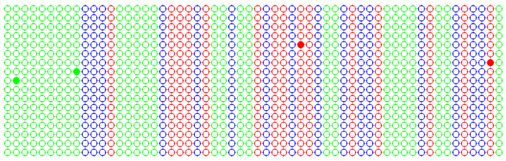 | CG (254) | **0.78%** |
| | | CHG (271) | **0%** |
| | | CHH (453) | **0.44%** |
| | | All (978) | **0.41%** |
| *B' mop3-1* sample 2 | 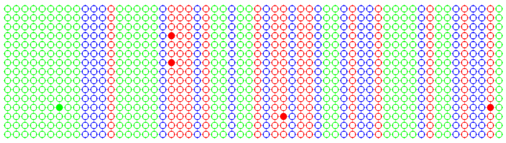 | CG (224) | **1.78%** |
| | | CHG (238) | **0%** |
| | | CHH (372) | **0.26%** |
| | | All (834) | **0.60%** |

●○ CG ●○ CHG ●○ CHH

**Supplemental Figure S8.** Targeted bisulfite data for *B'*, *B-I*, *B' mop1-1*, *B' Mop2/mop2-1* and *B' mop3-1* DNA derived from leaf 4 of V4 stage plants. DNA methylation profiles of A) The *B'* repeat junction region, and B) a *Fie2* sequence as a control for complete conversion are shown (Gutierrez-Marcos *et al.* 2006). Methylated and unmethylated cytosines are represented by filled and empty circles, respectively. The color code indicates the sequence context of individual cytosines. For each sample, the total number of cytosines and the fraction of non-converted cytosines in each sequence context are indicated.

### Supplemental Figure S9

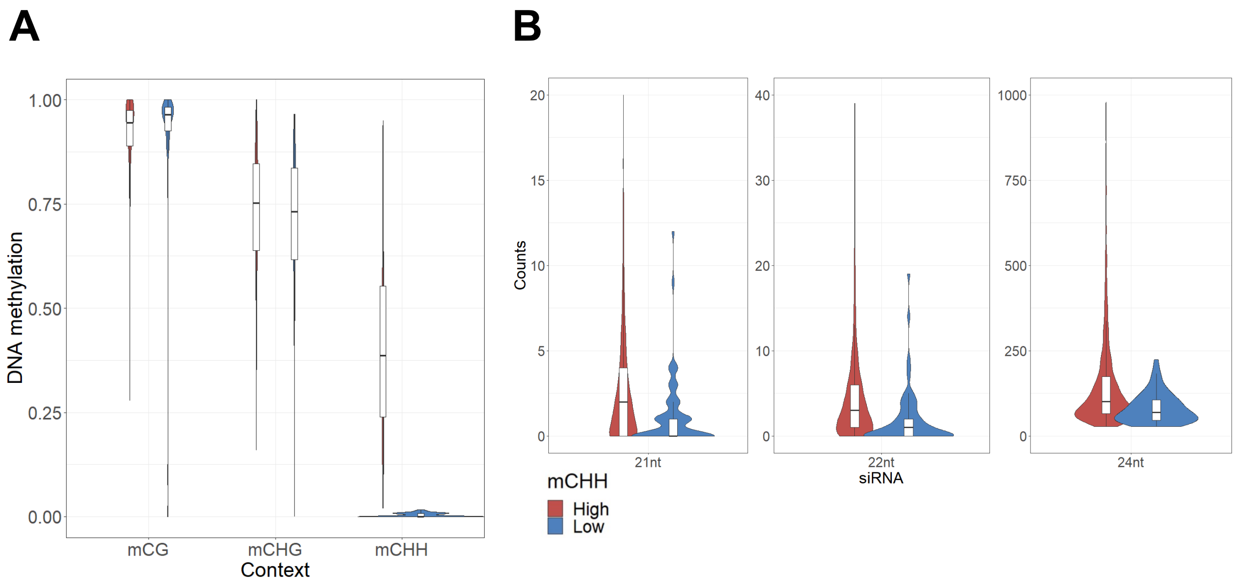

**Supplemental Figure S9.** Identification of low and high mCHH *mop1* loci and their respective siRNA levels. A) Violin plot showing DNA methylation levels (mC/total C) for low and high mCHH *mop1* loci. Data are from B73 developing ear (Gent et al., 2014), the source data used in defining low or high mCHH loci. B) Counts of uniquely-mapping siRNAs that overlapped at least half their lengths with low or high mCHH *mop1* loci. Data are from developing ear closely related to B73, the source of the *Mop1* wild-type siRNAs used in defining *mop1* loci (Gent *et al.*., 2014).

#### Supplemental Figure S10

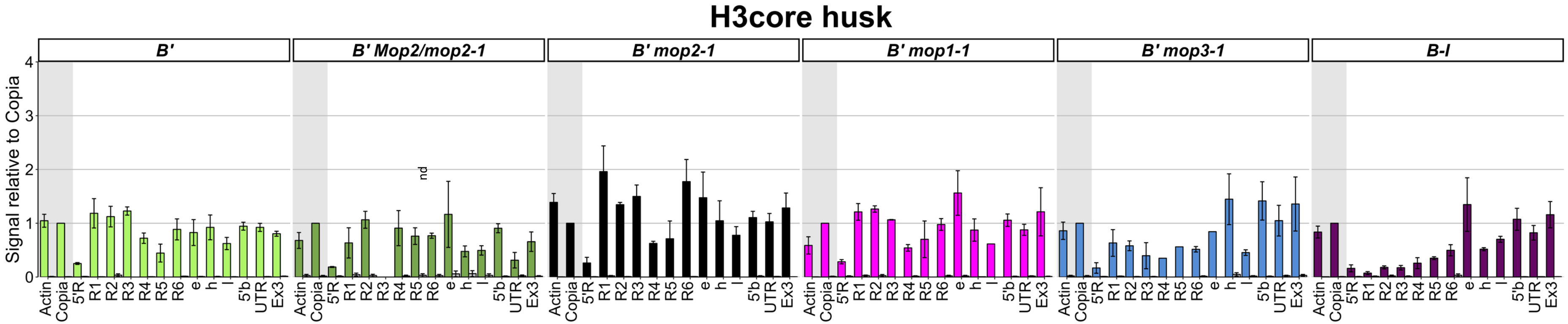

**Supplemental Figure S10.** No decrease in nucleosome occupancy was observed in the *mop* mutants. ChIP experiments were performed with an antibody recognizing histone H3, using husk tissue. ChIP signals were normalized to *Copia* signals. Colored bars indicate the ChIP signals, grey bars the no-antibody immunoprecipitations. The error bars indicate the SEM of two (*B' mop1-1*), three *(B'*, *B' Mop/mop2-1, B' mop3-1*) or four (*B' mop2-1*, *B-I*) replicate experiments.

### Supplemental Figure S11

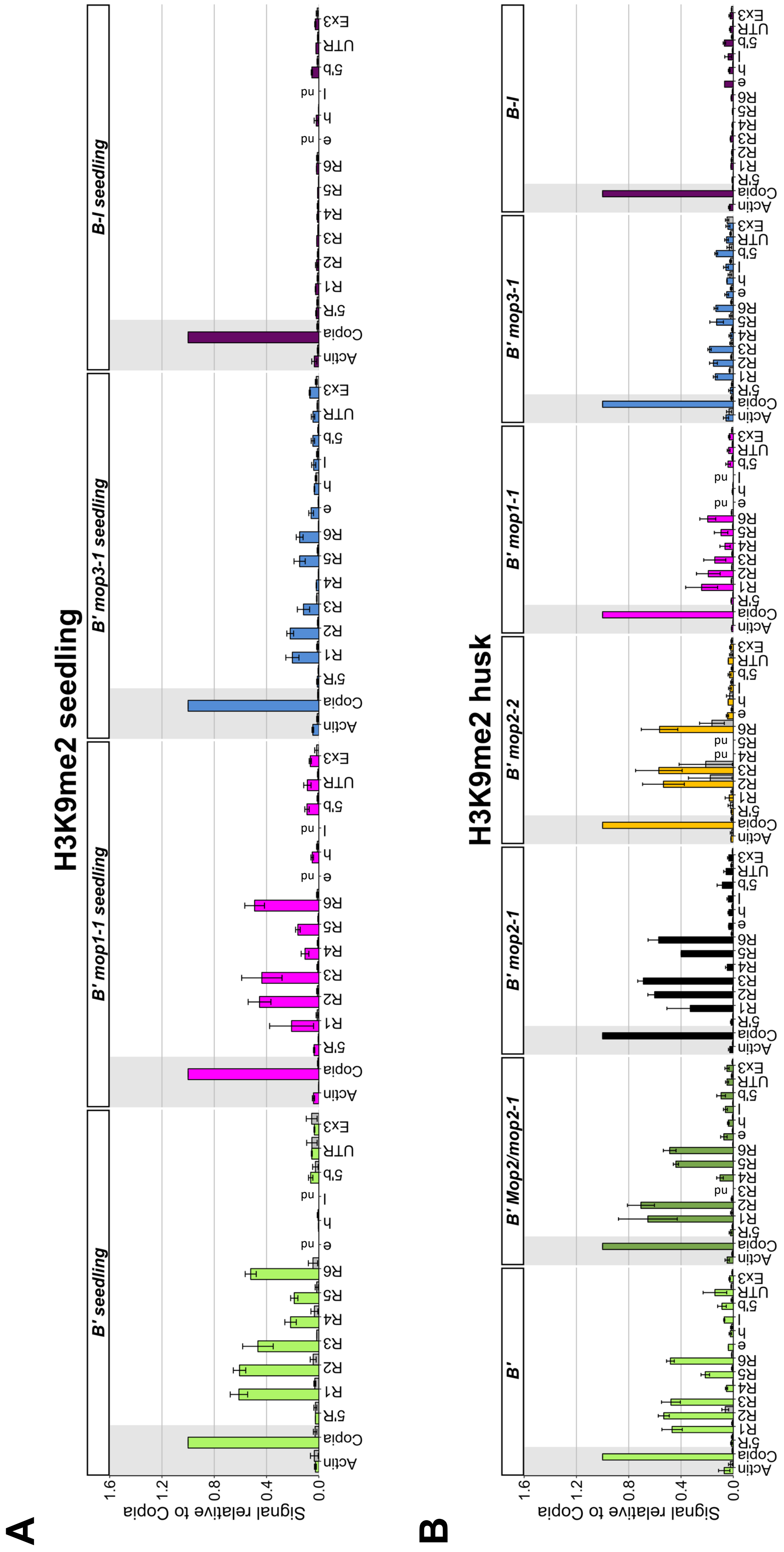

C

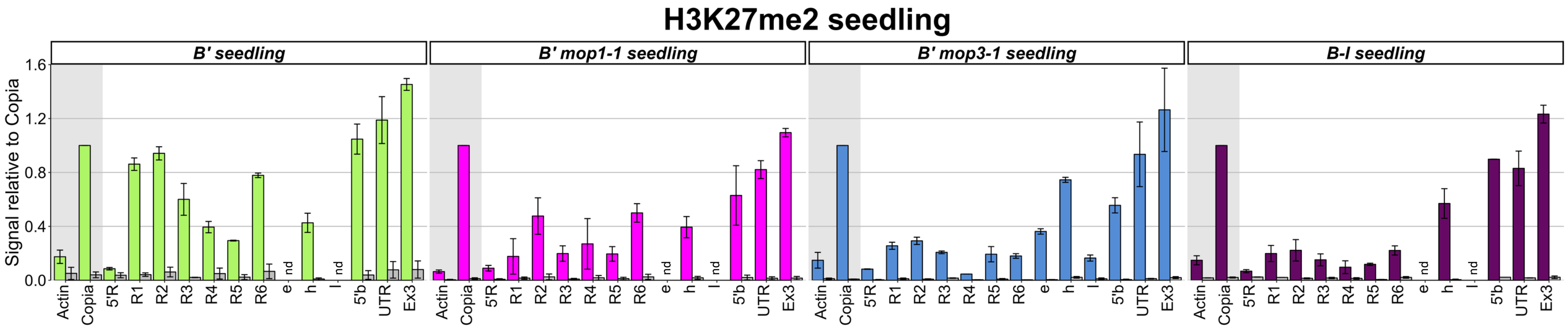

D

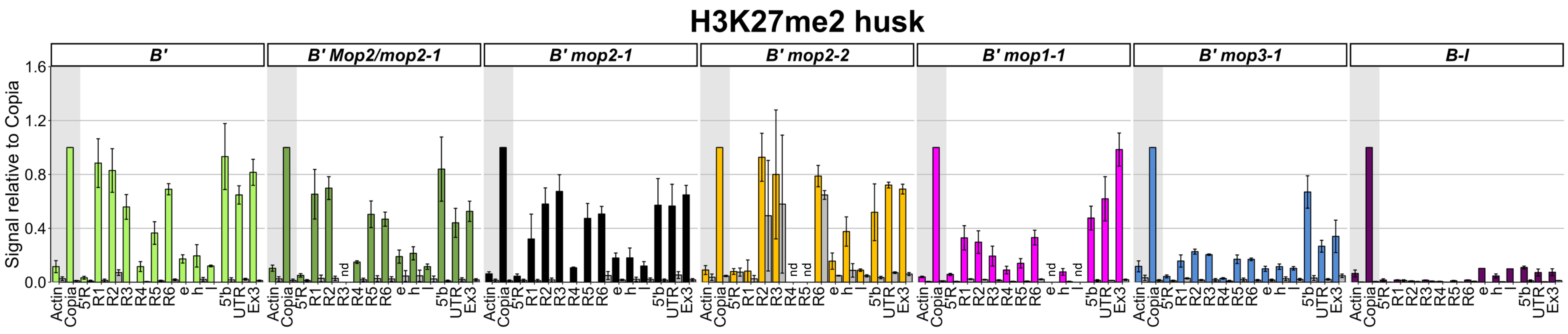

**Supplemental Figure S11.** Transcriptional activation in *mop1-1* and *mop3-1* is associated with reduced levels of repressive histone marks. ChIP-qPCR experiments were performed on seedling (A, C) and husk tissue (B, D) from *B'*, *B' Mop/mop2-1*, *B' mop2-1*, *B' mop2-2*, *B' mop1-1*, *B' mop3-1* and *B-I* plants with antibodies recognizing H3K9me2 (A, B), and H3K27me2 (C, D). Colored bars indicate the ChIP signals, grey bars the no-antibody control. ChIP signals were normalized to *Copia* signals. Error bars indicate the SEM of multiple biological replicates. For seedling tissue (A, C), three (*B', B' mop1-1*) or two (*B' mop3-1, B-I)* replicates were performed. For husks (B, D), three (*B', B' mop3-1*, *B-I*), four (*B' mop1-1, B' Mop/mop2-1*, *B' mop2-1)*, or two (*B' mop2-2*) replicates were performed. nd = no data for primer pair.
